# Upregulation of FasII underlies synergistic neuropathological and behavioral defects in a *Drosophila* model of myotonic dystrophy

**DOI:** 10.1101/2024.05.26.595976

**Authors:** Alex Chun Koon, Ka Yee Winnie Yeung, Lok I Leong, Zhefan Stephen Chen, Shaohong Isaac Peng, Joyce Man See Fung, Yitao Wu, Noah S. Armstrong, Ariadna Bargiela, Nerea Moreno, Javier Poyatos, Juan Vilchez, Paul Magneron, Aline Huguet, Cassandra Kussius Brewer, Erin Savner Beck, Rubén Artero, Mário Gomes-Pereira, Genevieve Gourdon, Vivian Budnik, C. Andrew Frank, Brian D. McCabe, Ho Yin Edwin Chan

## Abstract

Myotonic dystrophy type 1 (DM1) is a multisystemic disorder that has been extensively studied for decades, yet our understanding of its neuropathological aspect remains rudimentary. In this study, we characterized a novel model of DM1 neuropathology by expressing untranslated expanded *CUG* repeats at the *Drosophila* larval neuromuscular junction. In this model, both pre- and postsynaptic expression of *CUG* repeats participate to induce reduction of synaptic boutons, increase of arbor disassembly and impairment of larval locomotor activity. We found that the expression of *CUG* repeats caused an upregulation of the cell adhesion molecule, FasII (NCAM1 in mammals), in both the motor neurons and the body wall muscles. Knockdown of *fasII* was sufficient to rescue bouton numbers and locomotor impairment in this model. Further analyses identified the upregulation of the FasII-C isoform as a major contributor of these phenotypes. Remarkably, overexpressing the FasII-A-PEST+ isoform rescued the synaptic and behavioral defects, likely by outcompeting the upregulated FasII-C. Our study provided the foundation for a basic mechanism of synapse dysregulation in DM1.

## Introduction

Myotonic dystrophy type 1 (DM1), also known as Steinert disease, is the most common form of muscular dystrophy that begins in adulthood, affecting as many as one in every 2,100 individuals ^1^. DM1 is a multisystemic disorder characterized not only by progressive myotonia and muscle degeneration, but also presence of cataract, heart dysfunction and neuropathology ^2^. Congenital DM1 can also occur in infants and children who present with severe symptoms of muscle weakness and hypotonia rather than myotonia, as well as cognitive impairment ^3^. DM1 is caused by a CTG trinucleotide repeat expansion in the 3’ UTR of the *Dystrophia Myotonica Protein Kinase* (*DMPK*) gene ^4, 5, 6^. Normal individuals may have fewer than 37 CTG repeats, whereas DM1 patients may have hundreds or even thousands of repeats ^7^. The gain-of-function from the mRNA transcripts harboring these untranslated expanded *CUG* repeats has been shown to contribute to some of the major pathological features of DM1 independent of the *DMPK* locus ^8^. In fact, the severity of the disease correlates with the amount of the repeats and the age of onset ^9^. It has been demonstrated that the *CUG* repeat-containing transcripts are retained in the nucleus and recruited into ribonuclear foci ^10^. This in turn sequesters RNA-binding proteins, such as muscleblind-like (MBNL1), which in turn compromises the RNA-splicing machinery ^11, 12^.

RNA toxicity is well-known to be associated with the neurodegenerative features of numerous repeat expansion diseases, including polyglutamine diseases, spinocerebellar ataxias and C9ORF72-associated amyotrophic lateral sclerosis/frontotemporal dementia ^13^. DM1 is not an exception. Despite originally being coined as a muscular dystrophy, DM1 is known to manifest many different neuropathological features ^14, 15, 16^. DM1 patients commonly exhibit degeneration of the pigmentary retina and loss of photoreceptor neurons ^2^. In addition, they may exhibit cognitive impairment, speech and language difficulties, attention deficit, autism spectrum disorder, autistic features, sleep disorder, social anxiety and peripheral neuropathy ^17, 18, 19, 20, 21, 22, 23^. Nevertheless, comparing to the muscle pathology of DM1, the neuropathological aspect of DM1 is much less explored. It is known that, in neurons, mutant mRNA containing *CUG* repeats accumulate in ribonuclear foci within the nuclei and co-localize with MBNL1, similar to that in muscles ^12^. However, in a transgenic mouse model of DM1 (DM300), the expansion of 300 *CUG* repeats does not induce detectable neuropathology ^24^. By contrast, the expansion of 1,300 repeats in DMSXL mice results in motor neuropathy, suggesting that large *CUG* expansion can indeed result in neuropathology ^25^. At the neuromuscular junction of these DMSXL mice, the end-plates have decreased size and complexity, and 23% of the end-plates are disconnected from the axonal branches, and the motor neurons have defects in conducting action potentials ^25^. Moreover, synaptic proteins, such as RAB3A and synapsin I, are upregulated and hyperphosphorylated respectively in DMSXL mice, in association with synaptic transmission and behavioral deficits ^15^. Neuronal progeny cells derived from human embryonic stem cells carrying the DM1 mutation exhibit defects in neurite outgrowth and synapse formation ^16^. Despite these findings, the underlying mechanism of how expanded *CUG* repeats influences neurons and synapses remains unclear.

Here, we capitalized the highly stereotypical and accessible *Drosophila* larval neuromuscular junction (NMJ) system to study the effect of untranslated *CUG* repeats on synaptic functions and behavior. We found that the simultaneous expression of expanded *CUG* in the presynaptic motor neurons (MNs) and the postsynaptic body wall muscles (BWMs) results in synergistic functional and structural impairment of the NMJ. We characterized this DM1 model of neuropathology, and identified the upregulation of Fasciclin II (FasII) (the *Drosophila* orthologue of mammalian neural cell adhesion molecule 1, NCAM1) as a major contributor to the NMJ phenotypes we observed. Dysregulation of NCAM1 was also observed in the DMSXL mouse model and in DM1 patients, supporting the relevance of our findings in *Drosophila*. Overexpression of FasII-C, an isoform of FasII with no cytoplasmic domain, mimicked the phenotypes of expanded *CUG* expression at the NMJ. We were able to rescue our DM1 model by either knocking down the upregulated *fasII* or by overexpressing the major neuronal isoform of FasII at the NMJ, suggesting that we could potentially alleviate the loss of synapses by restoring the proper ratios of cell adhesion molecule isoforms. These findings provided significant new insights into the underlying mechanisms of DM1 neuropathology.

## Results

### Simultaneous pre- and postsynaptic expression of expanded *CUG* repeats causes neuromuscular defects

We used the *Drosophila* larval NMJ system to examine how *CUG* repeated-mediated toxicity affects the growth and development of neurons. The larval NMJ is characterized by its high level of plasticity during larval development and large synapses that are easy to visualize and to record from ^26^. Importantly, it also allows easy manipulation of the presynaptic MNs and the postsynaptic BWMs separately. *C380-Gal4 (C380)* is a presynaptic MN driver line, while *C57-Gal4 (C57)* is a postsynaptic BWM driver line. Using these Gal4 drivers, we overexpressed either the control construct (*UAS-CUG_60_*) (with the number of *CUG* repeats lower than the disease threshold) or the expanded *CUG* repeat construct (*UAS-CUG_480_*), which had been previous characterized in an adult *Drosophila* DM1 model ^27^. Similar to the *CUG_60_* control, we found that expression of *CUG_480_* with either *C380* or *C57* alone did not result in any change in synaptic bouton numbers in late 3^rd^ instar larvae. However, a robust bouton reduction phenotype was observed when both drivers were used to drive *CUG_480_* expression in both pre- and postsynaptic compartments simultaneously (Fig. 1a and 1b). This phenotype was not observed when we overexpressed another untranslated repeat expansion construct (*CAG_250_*) using both drivers, suggesting that the phenotype is specific to expanded *CUG* repeats (Fig. 1b).

**Figure 1.**
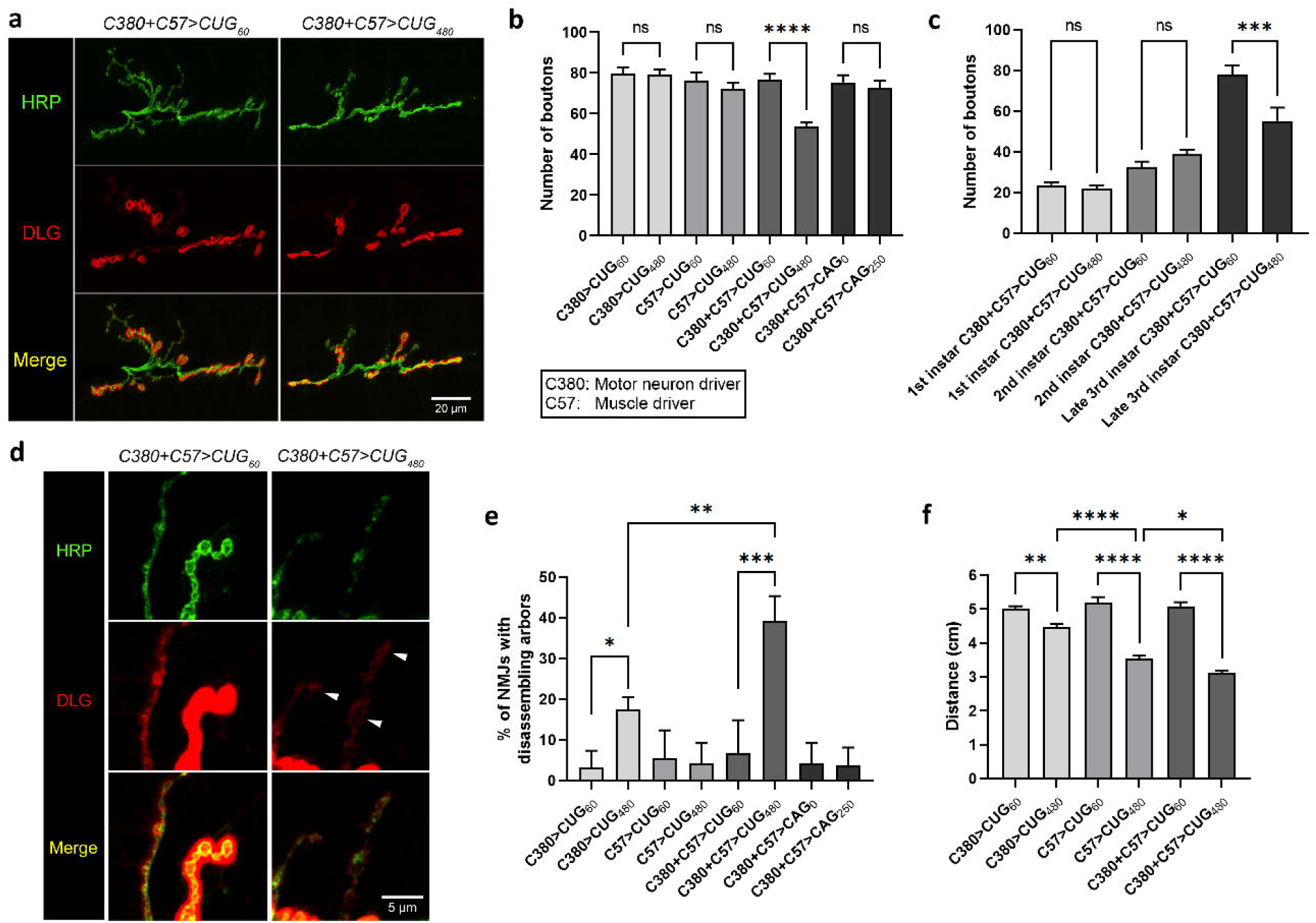
Simultaneous pre- and postsynaptic overexpression of *CUG_480_* causes NMJ and locomotor defects. **a** Confocal micrographs of *Drosophila* NMJs of late 3^rd^ instar larvae on muscles 6 and 7 of segment A3. Anti-HRP (in green) marks the presynaptic boutons. Anti-Discs large (DLG) (in red) marks the postsynaptic density. Scale bar is 20 μm. **b** Quantification of bouton numbers in **a** and in other related genotypes. Untranslated *CAG_0_*and *CAG_250_* were used as additional controls to show sequence specificity of the phenotype observed. *n* = 18, 17, 15, 19, 18, 39, 14, 16, where *n* is the number of analyzed NMJs. **c** Quantification of bouton numbers in 1^st^, 2^nd^ and late 3^rd^ instar larvae overexpressing either *CUG_60_* or *CUG_480_* using *C380* and *C57*. *n* = 16, 18, 14, 14, 10, 10, where *n* is the number of analyzed NMJs. **d** Confocal micrographs at high magnification showing arbors of boutons of NMJs at muscles 6 and 7 of segment A3. White arrowheads denote signs of disassembling boutons and arbors. Scale bar is 5 μm. **e** Quantification of disassembling arbors in **d** and in other related genotypes. *n* = 18, 17, 15, 19, 18, 39, 14, 16, where *n* is the number of analyzed NMJs. **f** Quantification of larval locomotor activity. *n* = 20, 20, 20, 20, 20, 30, where *n* indicates the number of analyzed larvae. Histograms depicts mean ± SEM. \**p* < 0.05, \*\**p* < 0.01, \*\*\**p* < 0.001, \*\*\*\**p* < 0.0001.

A phenotype of fewer boutons could have been caused by either reduced synaptic growth during development or disassembly/retraction of mature boutons after they were formed, or a combination of both. If the phenotype is present in the early stages of the larva, then it is likely due to reduced synaptic growth during development. Thus, we dissected 1^st^ instar and 2^nd^ instar larvae expressing *CUG_480_* using *C380* and *C57* drivers to investigate whether they had reduced bouton numbers. Interestingly, we found that both 1^st^ and 2^nd^ instar larvae had no detectable reduction of bouton numbers, while 3^rd^ instar had a robust decrease (Fig. 1c). These results suggest that the NMJs of *CUG_480_*-expressing animals, at least in terms of morphology, were rather normal during their early larval stages, and that the bouton reduction phenotype began only around the 3^rd^ instar stage.

A more direct way to determine whether synaptic disassembly/retraction is present in *CUG_480_*-expressing larvae would be to check for the presence of mature postsynaptic markers after the disassembly/retraction occurred. Discs large (DLG) is the *Drosophila* counterpart of mammalian postsynaptic MAGUK scaffolding proteins, including SAP-97, SAP-70 and PSD-95 ^28, 29^, and it is only present in the postsynaptic density of a mature bouton. Detailed analysis of the NMJs allowed us to identify some disconnected, disassembling boutons from seemingly degenerating arbors in larvae that expressed *CUG_480_* using both *C380* and *C57* drivers (Fig. 1d and 1e). We were able to observe the faint presence of DLG on the BWMs near these disassembling boutons, indicating that these were once mature boutons with postsynaptic densities (Fig. 1d). Therefore, although we cannot rule out the possibility that there might be reduced synaptic growth during development that contributed to the bouton reduction phenotype in these animals, we are certain that the disassembly of mature boutons and arbors, at least in part, contributed to this phenotype. Interestingly, although the expression of *CUG_480_* using the presynaptic MN driver (*C380*) alone did not give rise to a detectable bouton reduction phenotype (Fig. 1b), it did result in a small but significant increase of arbor disassembly (Fig. 1e). But the expression of *CUG_480_* using both presynaptic (*C380*) and postsynaptic (*C57*) drivers resulted in a significantly higher amount of arbor disassembly than using *C380* alone (Fig. 1e). These results suggest that although the expression of *CUG_480_* presynaptically alone does not yield a phenotype in bouton numbers, it does cause morphological defects. Also, our results suggest a synergistic effect on arbor disassembly when *CUG* RNA toxicity is present in both the presynaptic MNs and the postsynaptic BWMs.

Usually, when structural defects are present at the NMJ, functional defects are also present. Thus, after analyzing the morphology of the NMJs, we sought to determine if the expression of *CUG_480_* would affect larval crawling behavior as well. Previous studies suggested that the expression of *CUG_480_* in the postsynaptic BWMs alone was sufficient to produce locomotor defects in *Drosophila* larvae ^30^. Since the expression of *CUG_480_*in MNs alone was sufficient to cause arbor disassembly (Fig. 1e), we hypothesized that this may also be sufficient to cause locomotor defects in larvae. Indeed, we found that expression of *CUG_480_* using *C380* alone was sufficient to produce a small but significant locomotor defect in 3^rd^ instar larvae; expression of *CUG_480_* using the postsynaptic driver (*C57*) alone was sufficient to produce a stronger locomotor defect; and the expression of *CUG_480_* using both *C380* and *C57* drivers produced the strongest phenotype (Fig. 1f). These results suggest that *CUG* RNA toxicity in both the presynaptic MNs and the postsynaptic BWM contribute to a robust behavioral defect in the larva.

Summarizing the above results, we found that the simultaneous presynaptic and postsynaptic expression of expanded *CUG* repeats synergistically result in morphological defects at the NMJ, which are accompanied by locomotor behavioral defects. These results contribute to the establishment of a novel *Drosophila* model of DM1 neuropathology.

### FasII/NCAM is upregulated in DM1 models and DM1 patients

At this point, we sought to determine the cause of the phenotypes in our DM1 model. Although there are many molecules that can alter bouton numbers at the NMJ, not many are present in both the presynaptic MNs and postsynaptic BWMs, and regulate bouton numbers in a synergistic manner. One molecule with such potentials is the cell adhesion molecule FasII, the *Drosophila* orthologue of mammalian NCAM1. The regulation of bouton numbers by FasII at the NMJ is complex. First, overexpression of total FasII presynaptically, postsynaptically or both can lead to different phenotypes in bouton numbers ^31^. Second, *fasII* null mutants’ lethality can only be rescued when FasII is overexpressed in both the central nervous system (CNS) and BWMs ^32, 33^, indicating its importance in both the pre- and postsynapse. Most importantly, *fasII* hypomorphic mutants exhibit a synaptic retraction phenotype ^32, 33^, which we had observed in our DM1 NMJ model. Therefore, we hypothesized that FasII could be dysregulated by the expanded *CUG* RNA.

To test this hypothesis, we had to express *CUG_480_* in the CNS and muscles, and collect the respective tissues for mRNA or protein analysis. The MN driver *C380* was not suitable for this purpose, as its expression was limited to only a small number of neurons in the CNS. Thus, we utilized *elav^GeneSwitch^-Gal4* (*elav^GS^*), which is an inducible pan-neuronal Gal4 driver that will only be expressed under the presence of RU486 ^34^. This GeneSwitch driver was used to avoid expression of *CUG_480_* in the entire CNS too early during embryonic development, since expression of *CUG_480_*in CNS had been shown to affect viability ^27^. We expressed *CUG_480_* in the CNS and muscles of the larva using *elav^GS^* and *C57*, reared the animals in standard *Drosophila* medium containing 50 µM RU486 since hatching, dissected the CNS and muscle tissues at the 3^rd^ instar larval stage, and performed semi-quantitative RT-PCR to determine total *fasII* levels in these tissues. Our results demonstrated that *fasII* transcripts were indeed upregulated in both types of tissues (Fig. 2a-d).

**Figure 2.**
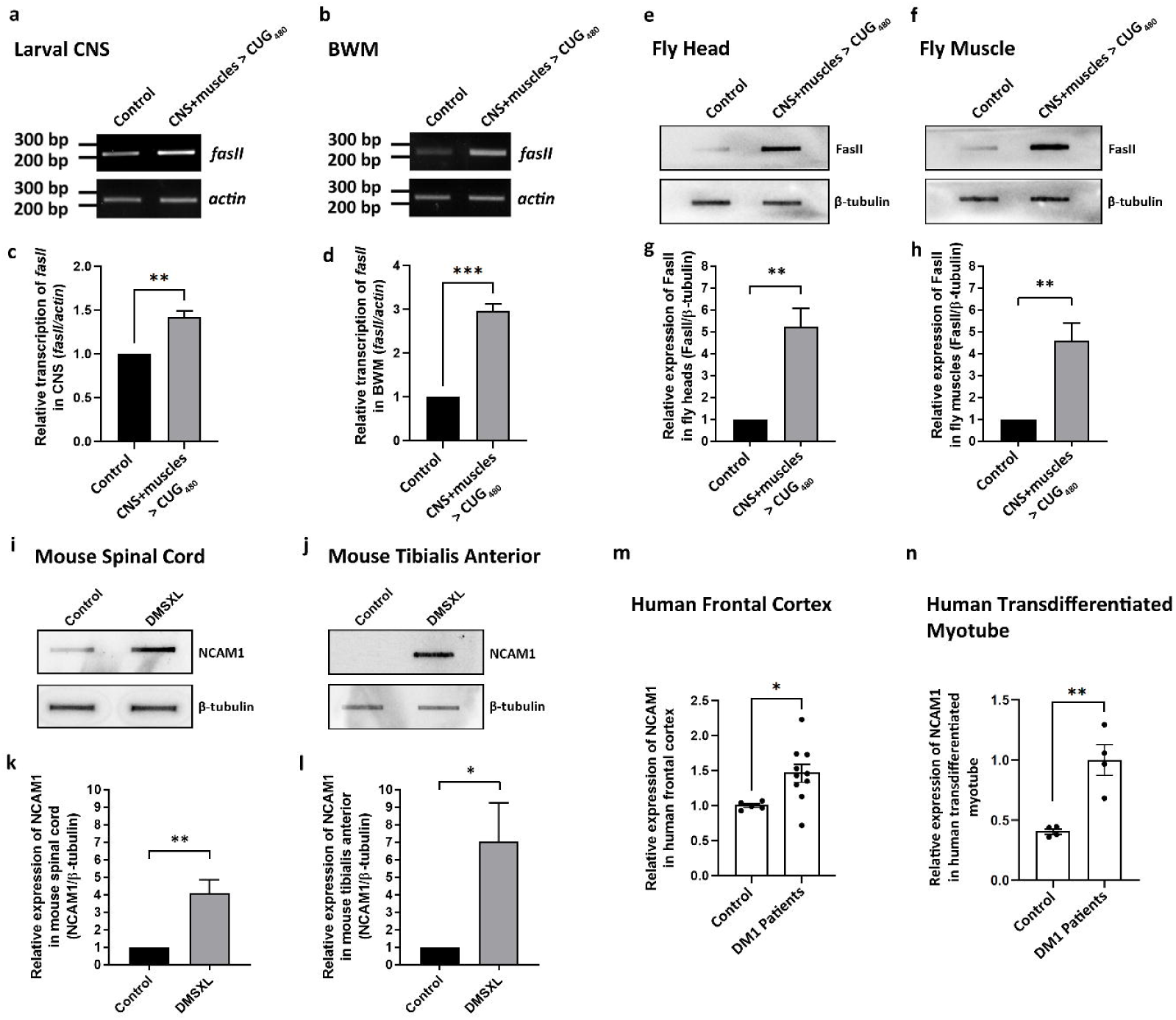
FasII/NCAM is upregulated in DM1 models and DM1 patients. **a, b** Representative semi-quantitative RT-PCR of *fasII* in *Drosophila* larval **(a)** CNS and **(b)** BWM. **c** Quantification of **a**. *n* = 3 for both genotypes. **d** Quantification of **b**. *n* = 3 for both genotypes. **e, f** Representative slot blots of FasII in adult *Drosophila* **(e)** head and **(f)** muscle. **g** Quantification of **e**. *n* = 3 for both genotypes. **h** Quantification of **f**. *n* = 3 for both genotypes. **i, j** Representative slot blots of NCAM1 in mouse **(i)** head and **(j)** muscle. **k** Quantification of **i**. *n* = 3 for both genotypes. **l** Quantification of **j**. *n* = 3 for both genotypes. **m** Quantified slot blot data of NCAM1 in human frontal cortex. *n* = 5 for control. *n* = 10 for DM1 patients. **n** Quantitative dot blot data of NCAM1 in human transdifferentiated myotube. *n* = 4 for both control and DM1 patients. Histograms depicts mean ± SEM. \**p* < 0.05, \*\**p* < 0.01, \*\*\**p* < 0.001.

Next, to show that this upregulation is not limited to just the larval stage, we let these animals growth until adulthood, collected the fly heads and muscles, and performed slot blot for total FasII proteins. Our results showed a significant upregulation of FasII in both the CNS and muscles of the *CUG_480_*-expressing adult flies (Fig. 2e-h). This demonstrated that not only the transcripts of *fasII* are upregulated by the expanded *CUG* repeats, but overall FasII proteins are also upregulated, and this upregulation lasts from the larval stages to adulthood.

To investigate whether such upregulation of cell adhesion molecules is present in a mammalian model of DM1, we analyzed neural and skeletal muscle samples from the DMSXL mice, which is a well-characterized mouse model of DM1 ^15, 35^. Using the slot blot technique, we found that NCAM1, the mammalian homolog of *Drosophila* FasII, was indeed upregulated in the spinal cord and the tibialis anterior of DMSXL mice (Fig. 2i-l), confirming that our findings in *Drosophila* is relevant to mammals.

To further verify that the upregulation of FasII in *Drosophila* and upregulation of NCAM1 in mice are relevant to actual DM1 patients, we performed slot blots on human NCAM1 using the frontal cortex of either control individuals or DM1 patients, and quantitative dot blots on transdifferentiated myotubes derived from either control or DM1 patient myoblasts. Once again, we found human NCAM1 to be upregulated in these types of tissues (Fig. 2m and 2n), confirming that our findings are truly relevant to DM1 patients.

### Knockdown of *fasII* rescues NMJs of *Drosophila* DM1 model

Since DM1 is known to be associated with splicing machinery defects ^11, 12^, we wondered if particular *fasII* isoforms were upregulated. There are at least four known isoforms of FasII in *Drosophila*. The two transmembrane isoforms of FasII (FasII-A-PEST+ and FasII-A-PEST-) are the major isoforms in neurons ^36^. FasII-C has no transmembrane domain and no cytoplasmic domain, and is tethered to the cell membrane via a GPI anchor ^37^. FasII-B is a predicted isoform that lacks a clear transmembrane domain and a GPI anchor, and is not well-characterized ^38^. To determine the transcript levels of the characterized *fasII* isoforms, we overexpressed *CUG_480_* using either *elav^GS^* or *C57*, and reared the animals in either medium containing 50 µM RU486 or plain medium, and dissected either the CNSs or BWMs at late 3^rd^ instar larval stage. Semi-quantitative RT-PCR was performed to analyze the transcript levels of *fasII-A-PEST+*, *fasII-A-PEST-* and *fasII-C* isoforms. To our surprise, all three *fasII* isoforms were found to be upregulated in both the CNS and BWMs (Fig. 3a and 3b).

**Figure 3.**
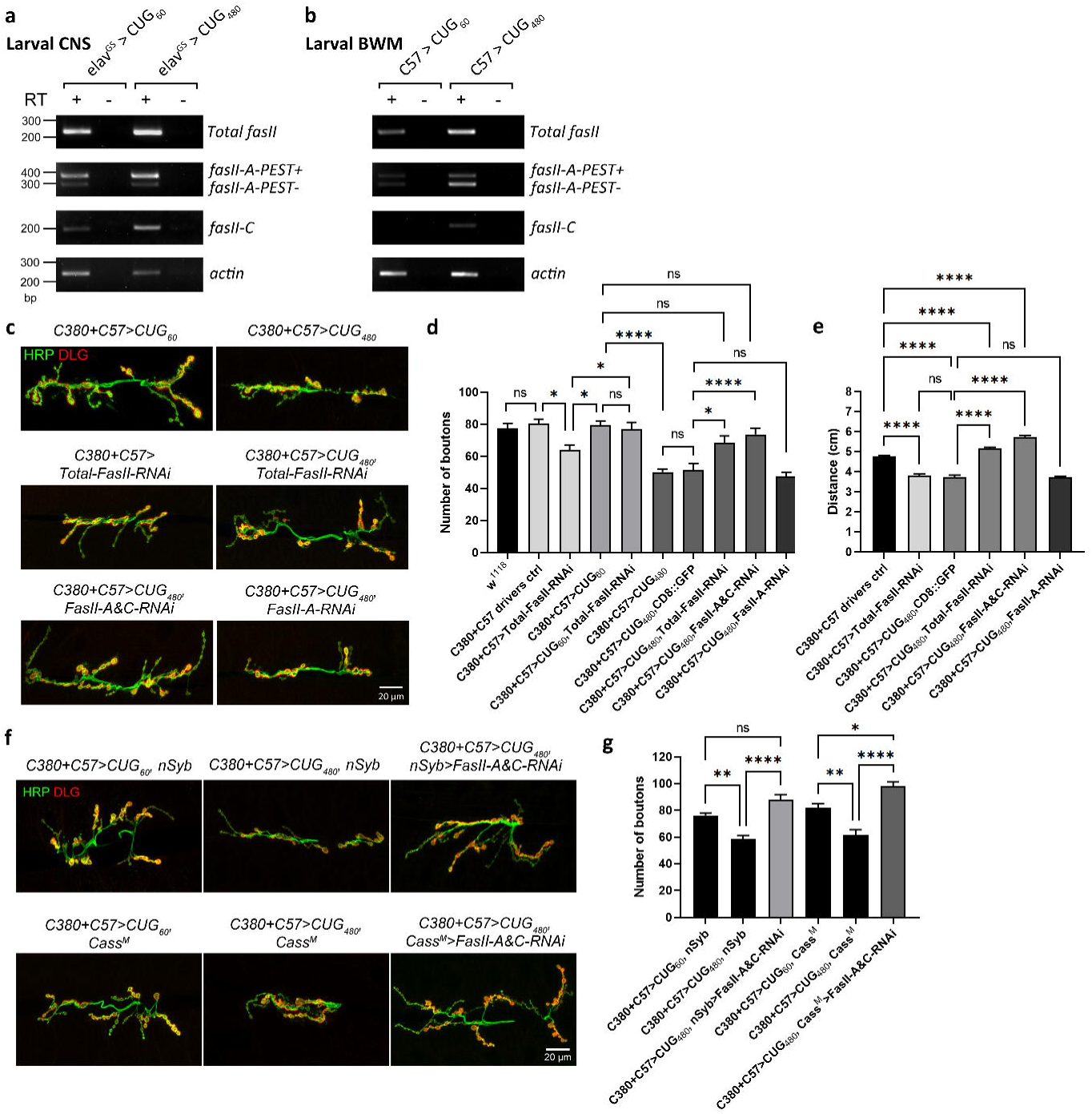
Knockdown of *fasII* rescues NMJs of *Drosophila* DM1 model. **a, b** Representative semi-quantitative RT-PCR of specific *fasII* isoforms in *Drosophila* larval **(a)** CNS and **(b)** BWM. **c** Confocal micrographs of *Drosophila* NMJs of late 3^rd^ instar larvae on muscles 6 and 7 of segment A3. Anti-HRP (in green) marks the presynaptic boutons. Anti-DLG) (in red) marks the postsynaptic density. Scale bar is 20 μm. **d** Quantification of bouton numbers in **c** and in other related genotypes. *n* = 12, 10, 13, 14, 12, 18, 13, 13, 19, 10, where *n* is the number of analyzed NMJs. **e** Quantification of larval locomotor activity. *n* = 40, 20, 18, 40, 20, 40, where *n* indicates the number of analyzed larvae. **f** Confocal micrographs of *Drosophila* NMJs of late 3^rd^ instar larvae on muscles 6 and 7 of segment A3. **g** Quantification of bouton numbers in **f**. *n* = 17, 14, 20, 15, 16, 20, where *n* is the number of analyzed NMJs. Histograms depicts mean ± SEM. \**p* < 0.05, \*\**p* < 0.01, \*\*\*\**p* < 0.0001.

If the NMJ morphological defects and larval crawling behavioral defects we observed in our DM1 model are caused by this upregulation of FasII in the MNs and BWMs, then we should be able to rescue these phenotypes by reducing the upregulated FasII in these tissues using RNAi. The fly line, *UAS-Total-FasII-RNAi*, had previously been characterized ^39, 40^. We first tested the effect of expressing *UAS-Total-FasII-RNAi* using the presynaptic *C380* and postsynaptic *C57* drivers in the control background, and found that it caused a small but significant decrease of synaptic bouton numbers (Fig. 3c and 3d). Next, we co-expressed *UAS-Total-FasII-RNAi* and *UAS-CUG_60_*. Interestingly, we found that the presence of *UAS-CUG_60_* nullified the bouton reduction effect caused by *UAS-Total-FasII-RNAi* (Fig. 3d). Next, we sought to determine if *UAS-Total-FasII-RNAi* can rescue the bouton reduction phenotype caused by *UAS-CUG_480_*. However, one caveat of the experiment is that the extra *UAS* from the RNAi construct might reduce the available Gal4 to express *UAS-CUG_480_*, which might result in a false rescue. Thus, we introduced a *UAS-CD8::GFP* to control for the extra *UAS* in the rescue genotype. Our results showed no significant difference between the bouton numbers of *C380+C57>CUG_480_*and *C380+C57>CUG_480_, CD8::GFP*, indicating that the extra *UAS* transgene expression did not significantly affect the bouton reduction phenotype caused by *UAS-CUG_480_* (Fig. 3d). We then co-expressed *UAS-Total-FasII-RNAi* and *UAS-CUG_480_*, and found that the knockdown of *fasII* indeed rescued the bouton numbers (Fig. 3c and 3d).

To further examine whether the knockdown of particular *fasII* isoforms were responsible for this rescue, we employed two fly lines that were designed to knockdown specific *fasII* isoforms. *UAS-FasII-A-RNAi* can knockdown both *fasII-A-PEST+* and *fasII-A-PEST-*, and had been previously characterized ^39^. *UAS-FasII-C-RNAi* was designed to target Exon 8 of *fasII*, which encodes for the GPI anchor. Thus, it was expected to knockdown *fasII-C* only. However, to our surprise, when we performed RT-PCR analysis on *Drosophila* BWMs expressing this construct, we found that it actually knocked down both *fasII-A* isoforms in addition to *fasII-C* (Supplementary Fig. 1 and Supplementary Fig. 2). Therefore, the line was renamed as *UAS-FasII-A&C-RNAi*. When we co-expressed this line and *UAS-CUG_480_*, we found that it rescued the bouton number, similar to *UAS-Total-FasII-RNAi* (Fig. 3c and 3d). In contrast, expression of *UAS-FasII-A-RNAi* was unable to rescue the bouton numbers (Fig. 3c and 3d), suggesting that the rescue in our DM1 model was not due to the knockdown of the *fasII-A* isoforms.

As we are able to rescue the bouton numbers at the NMJ with *fasII* knockdown, we further sought to determine if this manipulation can also rescue larval crawling behavior. Indeed, consistent to our findings regarding bouton numbers, we found that *UAS-Total-FasII-RNAi* and *UAS-FasII-A&C-RNAi* were capable of rescuing crawling behavior in the DM1 model background, while *FasII-A-RNAi* was unable to do so (Fig. 3e). Intriguingly, these constructs actually produced an over-rescue phenotype in locomotor activity.

In Fig. 1, we found that expression of *CUG_480_* in either the presynaptic MNs or the postsynaptic BWMs alone was not sufficient to cause reduction in bouton numbers. Thus, hypothetically, rescuing either the presynaptic MNs or the postsynaptic BWMs using *fasII* knockdown should be sufficient to rescue the bouton numbers at the NMJ in our DM1 model. To test this hypothesis, we needed to perform our tissue-specific *fasII* knockdown independent of the Gal4/UAS system since *CUG_480_* is being overexpressed by *C380* and *C57* simultaneously in our DM1 model. To achieve this, we employed the LexA/LexAop binary expression system in conjunction with the Gal4/UAS system ^41^. *nSyb-LexA (nSyb)* is a pan-neuronal driver line that expresses in all neurons, while *Cass^M^-LexA (Cass^M^)* is a muscle driver line that expresses in BWMs (Supplementary Fig. 3). We generated *LexAop-FasII-A&C-RNAi* using the same construct used in *UAS-FasII-A&C-RNAi*. Similar to *UAS-FasII-A&C-RNAi*, we found that *LexAop-FasII-A&C-RNAi* also knocks down both *fasII-A* and *fasII-C* isoforms (Supplementary Fig. 4). We then expressed *LexAop-FasII-A&C-RNAi* to knockdown *fasII* using either *nSyb* or *Cass^M^* in the DM1 model background of *C380+C57>CUG_480_*. As we expected, knockdown of *fasII-A* and *fasII-C* using either *nSyb* or *Cass^M^* was sufficient to rescue the bouton numbers in our DM1 model (Fig. 3f and 3g). In fact, knockdown of *fasII-A* and *fasII-C* using *Cass^M^*resulted in an over-rescue phenotype. These results suggest that the upregulation of FasII by *CUG_480_*in this DM1 model has to be present in both the presynaptic MNs and the postsynaptic BWMs to cause reduction of boutons at the NMJ.

### Overexpression of FasII-C results in NMJ morphological defects and behavioral defects

At this point, we had accumulated a number of observations. Firstly, a previous study suggested that FasII-C is usually not the major isoform in neurons ^36^, which we were able to confirm in our results (Fig. 3a). In addition, we found that FasII-C is also the least expressed isoform in wild-type BWMs (Fig. 3b). Thus, its upregulation in CUG_480_-expressing MNs and BWMs may negatively impact NMJ functions. Secondly, the fact that *UAS-Total-FasII-RNAi* and *UAS-FasII-A&C-RNAi*, but not *UAS-FasII-A-RNAi*, were able to rescue bouton numbers and behaviour (Fig. 3c and 3d), suggests that the upregulation of FasII-C may be the cause of the NMJ defects in our DM1 model. To test this hypothesis, we overexpressed *UAS-FasII-A-PEST+*, *UAS-FasII-A-PEST-* or *UAS-FasII-C* pre- and postsynaptically using *C380* and *C57* driver simultaneously (in the non-CUG_480_-expressing *w^1118^* control background). We found that overexpression of FasII-A-PEST+ or FasII-A-PEST-had no impact on bouton numbers, while overexpression of FasII-C caused a reduction in bouton numbers, similar to our DM1 model (Fig. 4a and 4b). Furthermore, overexpression of FasII-C also resulted in an increase of disassembling arbors (Fig. 4c and 4d), which is a phenotype we observed when we overexpressed *CUG_480_* at the NMJ (Fig. 1d and 1e).

**Figure 4.**
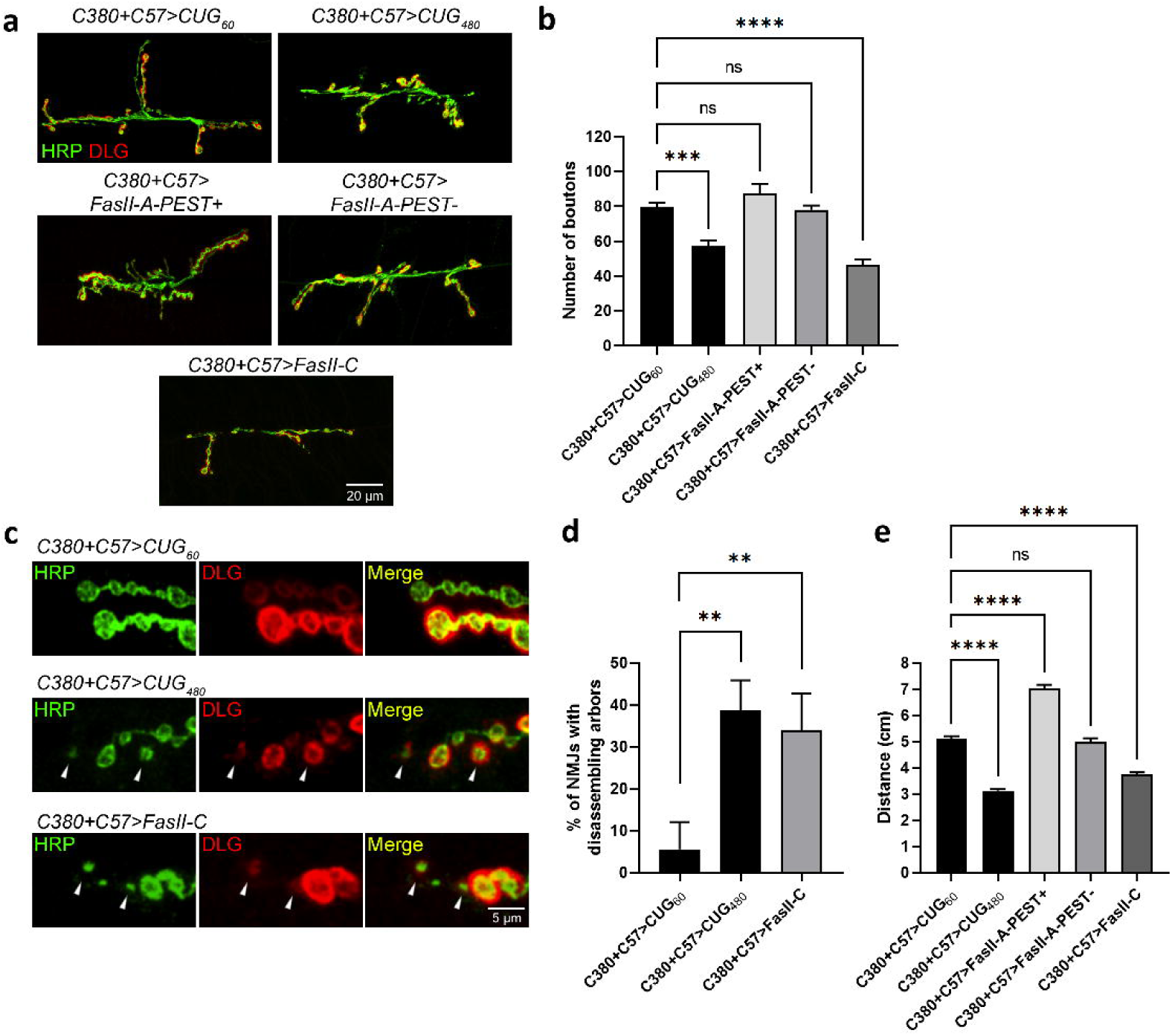
Overexpression of FasII-C causes in NMJ morphological defects and larval locomotor impairment. **a** Confocal micrographs of *Drosophila* NMJs of late 3^rd^ instar larvae on muscles 6 and 7 of segment A3. Anti-HRP (in green) marks the presynaptic boutons. Anti-DLG (in red) marks the postsynaptic density. Scale bar is 20 μm. **b** Quantification of bouton numbers in **a**. *n* = 14, 13, 13, 10, 11, where *n* is the number of analyzed NMJs. **c** Confocal micrographs at high magnification showing arbors of boutons of NMJs at muscles 6 and 7 of segment A3. White arrowheads denote signs of disassembling boutons and arbors. Scale bar is 5 μm. **d** Quantification of disassembling arbors in **c**. *n* = 12, 18, 16, where *n* is the number of analyzed NMJs. **e** Quantification of larval locomotor activity. *n* = 20, 20, 30, 20, 20, 20, where *n* indicates the number of analyzed larvae. Histograms depicts mean ± SEM. \**p* < 0.05, \*\**p* < 0.01, \*\*\**p* < 0.001, \*\*\*\**p* < 0.0001.

In terms of larval crawling behavior, overexpression of FasII-A-PEST+ pre- and postsynaptically resulted in a significant increase of locomotor activity; overexpression of FasII-A-PEST- had no impact; and overexpression of FasII-C resulted in a decrease of locomotor activity, which once again mimicked the phenotype in our DM1 model (Fig. 4e).

### Simultaneous pre- and postsynaptic overexpression of FasII-C synergistically causes functional defects at the NMJ

To further evaluate the impacts of FasII-C overexpression on NMJ functions, we performed electrophysiological analyses to examine synaptic transmission. We overexpressed *UAS-FasII-C* using either *C380*, *C57* or both drivers simultaneously, and measured the amplitude and frequency of mEPSPs and amplitudes of EPSPs. We found that overexpression of FasII-C presynaptically or postsynaptically alone had no impact on mEPSP amplitude (Fig. 5a and 5b). In contrast, simultaneous pre- and postsynaptic overexpression of FasII-C resulted in a significant decrease of mEPSP amplitude, indicating a decrease in vesicle size or a decrease in glutamate receptors at the NMJ (Fig. 5c). In terms of EPSP, both presynaptic and postsynaptic overexpression of FasII-*C* resulted in a small but significant decrease, while pre- and postsynaptic co-overexpression resulted in an apparent further decrease (Fig. 5d-f), indicating an overall decrease of evoked response. In terms of quantal content, presynaptic overexpression had no impact, postsynaptic overexpression resulted in a small but significant decrease, while simultaneous pre- and postsynaptic overexpression resulted in a further decrease (Fig. 5g-i), indicating less vesicles being released per evoked response. Lastly, in terms of mEPSP frequency, presynaptic overexpression resulted in small but significant decrease, postsynaptic overexpression had no impact, while simultaneous pre- and postsynaptic overexpression caused a further decrease (Fig. 5j-l), indicating fewer vesicles being spontaneously released. All these data strongly suggest that presynaptic and postsynaptic overexpression of *FasII-C* synergistically causes functional defects to the NMJ.

**Figure 5.**
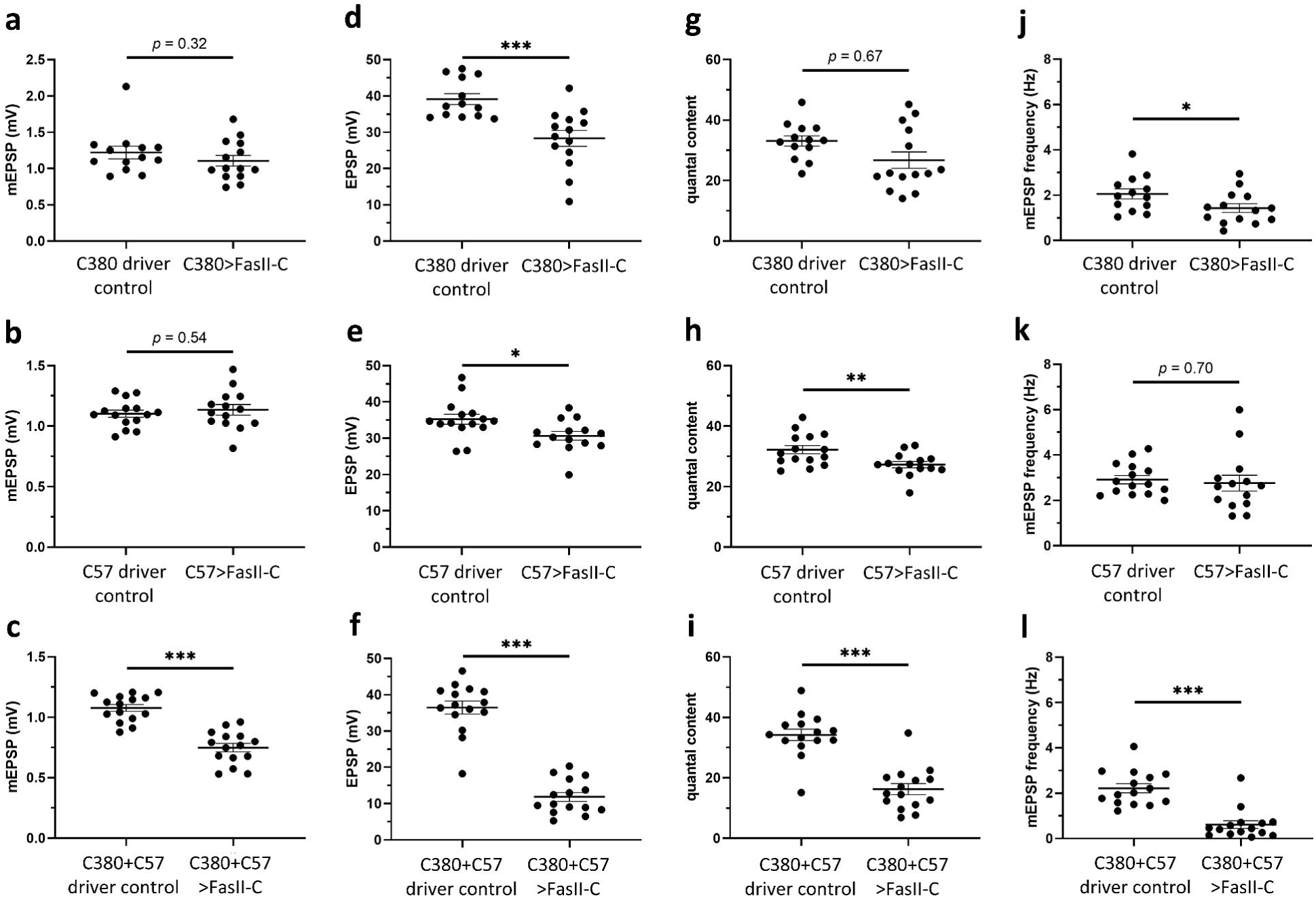
Simultaneous pre- and postsynaptic overexpression of FasII-C synergistically impairs synaptic functions at the *Drosophila* larval NMJ. **a-c** mEPSP amplitude of *Drosophila* larval NMJ overexpressing FasII-C using **(a)** *C380*, **(b)** *C57*, **(c)** *C380+C57*. **d-f** EPSP amplitude of *Drosophila* larval NMJ overexpressing FasII-C using **(d)** *C380*, **(e)** *C57*, **(f)** *C380+C57*. **g-i** Quantal content of *Drosophila* larval NMJ overexpressing FasII-C using **(g)** *C380*, **(h)** *C57*, **(i)** *C380+C57*. **j-l** mEPSP frequency of *Drosophila* larval NMJ overexpressing FasII-C using **(j)** *C380*, **(k)** *C57*, **(l)** *C380+C57*. Histograms depicts mean ± SEM. \**p* < 0.05, \*\**p* < 0.01, \*\*\**p* < 0.001.

### Overexpression of FasII-A rescues bouton numbers at the larval NMJs of the *Drosophila* DM1 model

Out of all three characterized isoforms of FasII, both FasII-A isoforms have a cytoplasmic domain that can potentially participate in intracellular signals, which may be important for maintaining synapse functionality. This cytoplasmic domain can also interact with DLG at the postsynapse ^42, 43^, which may facilitate retrograde signals required for synapse maintenance ^44^. FasII-C, however, lacks a cytoplasmic domain. Thus, in our DM1 model, if increased amounts of FasII-C are present at the synapse, they may compete with other FasII isoforms to bind with FasII-A, which may in turn disrupt the intracellular signals at the synapse. Therefore, we wondered if it is possible to restore the proper ratio of FasII isoforms in our DM1 model by overexpressing FasII-A. By outcompeting the upregulated FasII-C with more FasII-A, we may restore the intracellular signals and rescue the bouton defects. To test this hypothesis, we overexpressed either *UAS-FasII-A-PEST+* or *UAS-FasII-A-PEST-* in conjunction with *UAS-CUG_480_* using *C380* and *C57*. Indeed, we found that the overexpression of either FasII-A isoforms was capable of rescuing the bouton numbers in our DM1 model (Fig. 6a and 6b). Moreover, overexpression of *UAS-FasII-A-PEST+* was capable of rescuing the locomotor impairment in our DM1 model (Fig. 6c). In fact, it resulted in an over-rescue (Fig. 6c). However, overexpression of *UAS-FasII-A-PEST-* did not result in observable improvement on locomotor activity (Fig. 6c).

**Figure 6.**
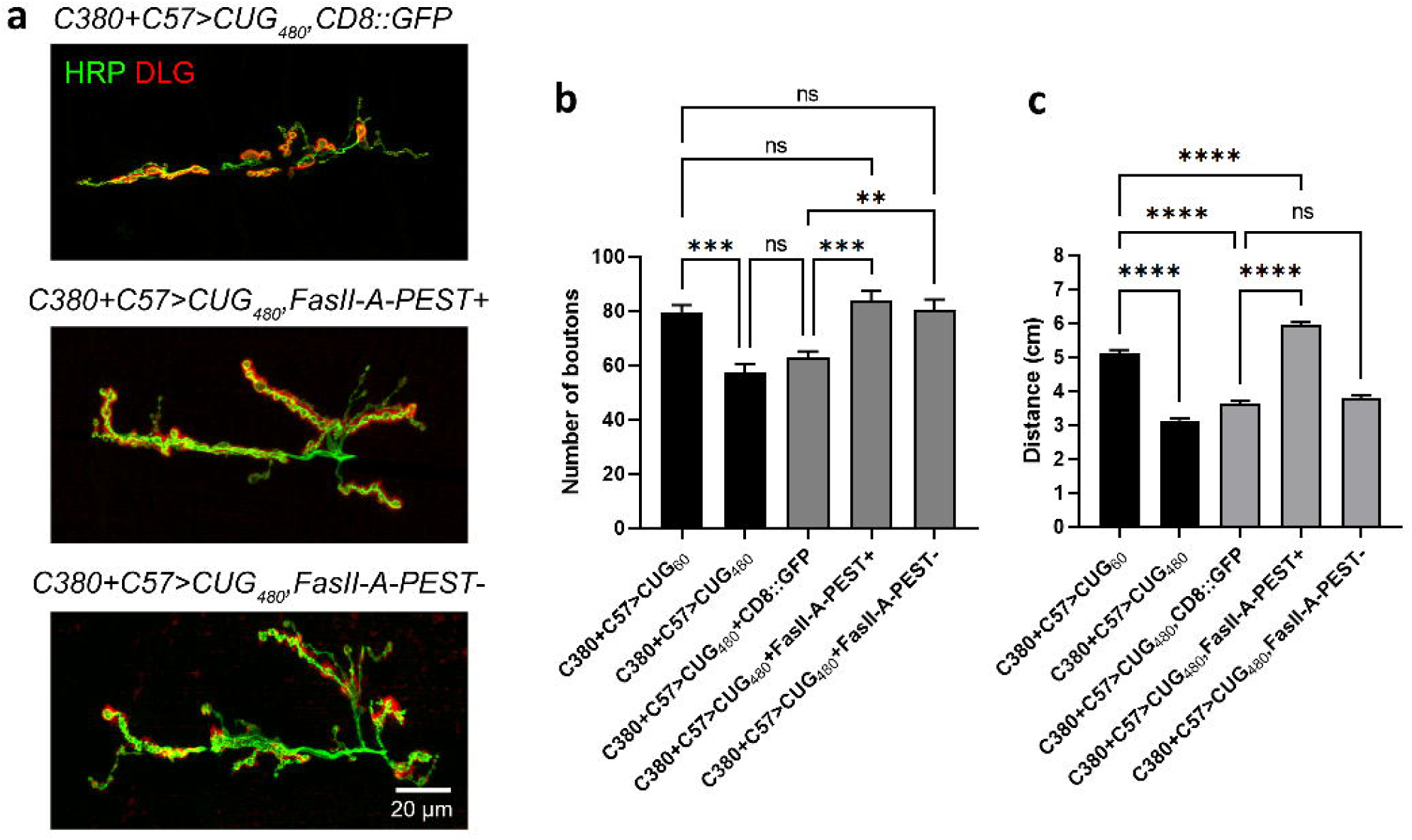
Overexpression of FasII-A rescues bouton numbers at the larval NMJs of the *Drosophila* DM1 model. **a** Confocal micrographs of *Drosophila* NMJs of late 3^rd^ instar larvae on muscles 6 and 7 of segment A3. Anti-HRP (in green) marks the presynaptic boutons. Anti-DLG (in red) marks the postsynaptic density. Scale bar is 20 μm. **b** Quantification of bouton numbers in **a**. *n* = 14, 13, 17, 20, 24, where *n* is the number of analyzed NMJs. **c** Quantification of larval locomotor activity. *n* = 20, 30, 20, 20, 20, where *n* indicates the number of analyzed larvae. Histograms depicts mean ± SEM. \*\**p* < 0.01, \*\*\**p* < 0.001, \*\*\*\**p* < 0.0001.

## Discussion

DM1 is widely recognized as a multisystemic disorder with neurological manifestations, including both peripheral nervous system and CNS abnormalities ^14^. Yet, despite decades of studies, the neuropathology in DM1 is still one of the most poorly understood aspects of the disease. In this study, we established a novel DM1 model of neuropathology using the *Drosophila* larval NMJ by expressing untranslated expanded *CUG* repeats pre- and postsynaptically. We observed a synaptic bouton reduction phenotype at the NMJ that only occurs when *CUG_480_* is simultaneously expressed in the presynaptic MNs and postsynaptic BWMs (Fig. 1). In addition, both pre- and postsynaptic expression of *CUG_480_* contributed to an arbor disassembly phenotype and larval locomotion impairment (Fig. 1). We determined that the expression of *CUG_480_* caused an upregulation of the cell adhesion molecule, FasII, at the NMJ (Fig. 2a-f). Similar upregulation of NCAM1 was also observed in a mouse model of DM1 and in DM1 patients (Fig. 2g-n). Remarkably, we were able to rescue the reduced bouton phenotype in our *Drosophila* DM1 model by knocking down total *fasII* either presynaptically or postsynaptically (Fig. 3f and 3g). We found that overexpression of FasII-C pre- and postsynaptically mimicked the NMJ morphological and behavioral phenotypes of our DM1 model (Fig. 4), as well as synergistically caused synaptic transmission defects (Fig. 5). Lastly, we demonstrated that overexpression of either FasII-A isoforms could rescue bouton numbers in our DM1 model (Fig. 6a and 6b), likely by outcompeting the upregulated FasII-C and restoring the proper ratio of FasII isoforms at the NMJ. One of the two FasII-A isoforms, FasII-A-PEST+, was also capable of rescuing the locomotor defects in our DM1 model (Fig. 6c).

After we found that total *fasII* knockdown was able to rescue our DM1 model, we originally intended to investigate the effect *fasII-C* knockdown using *UAS-FasII-C-RNAi*. This construct was generated by targeting Exon 8 of the *fasII* gene, which encodes for the GPI anchor. However, our results surprisingly showed that this construct actually knocks down the two *fasII-A* isoforms as well (Supplementary Fig. 1), which prompted us to rename it as *UAS-FasII-A&C-RNAi*. One possible explanation that we cannot exclude is the common off-target effect of RNAi, which somehow allowed the amplicon to bind to other exons. A second possibility is that the two characterized FasII-A isoforms actually also contain a GPI anchor in addition to having a transmembrane domain. A third possibility is that there are other uncharacterized FasII isoforms that contain a GPI anchor. A recent study indeed found a second GPI anchor-containing FasII isoform ^45^. However, the reverse RT-PCR primer for *fasII-A* in our study partially targets the nucleotide sequence of Exon 7 and the transmembrane domain (TM). This actually suggests that this second GPI-anchor isoform contains a TM domain, which differs from what Neuert et al. had found ^45^. Furthermore, when we used a pair of primers that exclusively amplifies *fasII-A-PEST+*, we still observed the knockdown by *UAS-FasII-A&C-RNAi* (Supplementary Fig. 2), which strongly indicates that the RNAi line can truly knockdown *fasII-A-PEST+*. Note that the two latter possible explanations above are not mutually exclusive, and they certainly warrant further investigations.

NCAM1 is the mammalian orthologue of *Drosophila* FasII. Both NCAM1 and FasII belong to the immunoglobulin (Ig) domain superfamily and possess homophilic cell-cell adhesion mediator activity ^46^. In humans, abnormal NCAM1-positive myofibers were observed in DM1 patient deltoid muscles ^47^, which might have been caused by an increased expression of NCAM1. *NCAM1* transcripts have also been reported to be upregulated in the frontal cortex of DM1 patients ^48^. In agreement with these previous reports, our data also showed increased levels of NCAM1 proteins in the spinal cord and tibialis anterior of DMSXL mice, as well as in the frontal cortex of DM1 patients and transdifferentiated myotubes derived from DM1 patient cells. However, we observed that the deltoids of DM1 patients actually had a lower expression NCAM1 than control individuals (Supplementary Fig. 5). It is possible that different types of skeletal muscles in higher organisms have different NCAM1 dysregulations under the toxicity of the expanded *CUG* repeats. Nevertheless, dysregulation of NCAM1 was still detected in human deltoids, which would likely result in synapse dysfunction at the neuromuscular junction. Alternatively, there could also be some kind of compensatory mechanisms in humans that occur in mature muscle cells, which may explain the high NCAM1 in transdifferentiated myotubes but low NCAM1 in deltoid muscles.

In our proposed model (Fig. 7), both FasII-A-PEST+ and FasII-A-PEST-are the major isoforms expressed at the presynaptic MNs and postsynaptic BWMs of a normal NMJ. These isoforms harbor a cytoplasmic domain, allowing them to convey intracellular signals. At the postsynaptic BWMs, the cytoplasmic PDZ-interaction motif of FasII-A isoforms interacts with DLG ^42, 43, 46^, which may facilitate retrograde signaling via molecules, such as Gbb (*Drosophila* counterpart of mammalian BMP), to maintain bouton numbers ^44^. Under the DM1-like condition, the expanded *CUG* repeats cause the upregulation of total FasII. The upregulated FasII-C may either bind to itself or compete with other isoforms to bind to the FasII-A isoforms via their Ig domains (the domains that mediate adhesion via transhomophilic binding). However, since FasII-C does not contain a cytoplasmic domain, it cannot transduce intracellular signaling like the FasII-A isoforms do ^46^. Hence, this binding of FasII-C to FasII-A disrupts the intracellular signals required for proper synaptic functions, thereby leading to defects in synaptic transmission, locomotion impairment, and even the gradual disassembly of the synapses.

**Figure 7.**
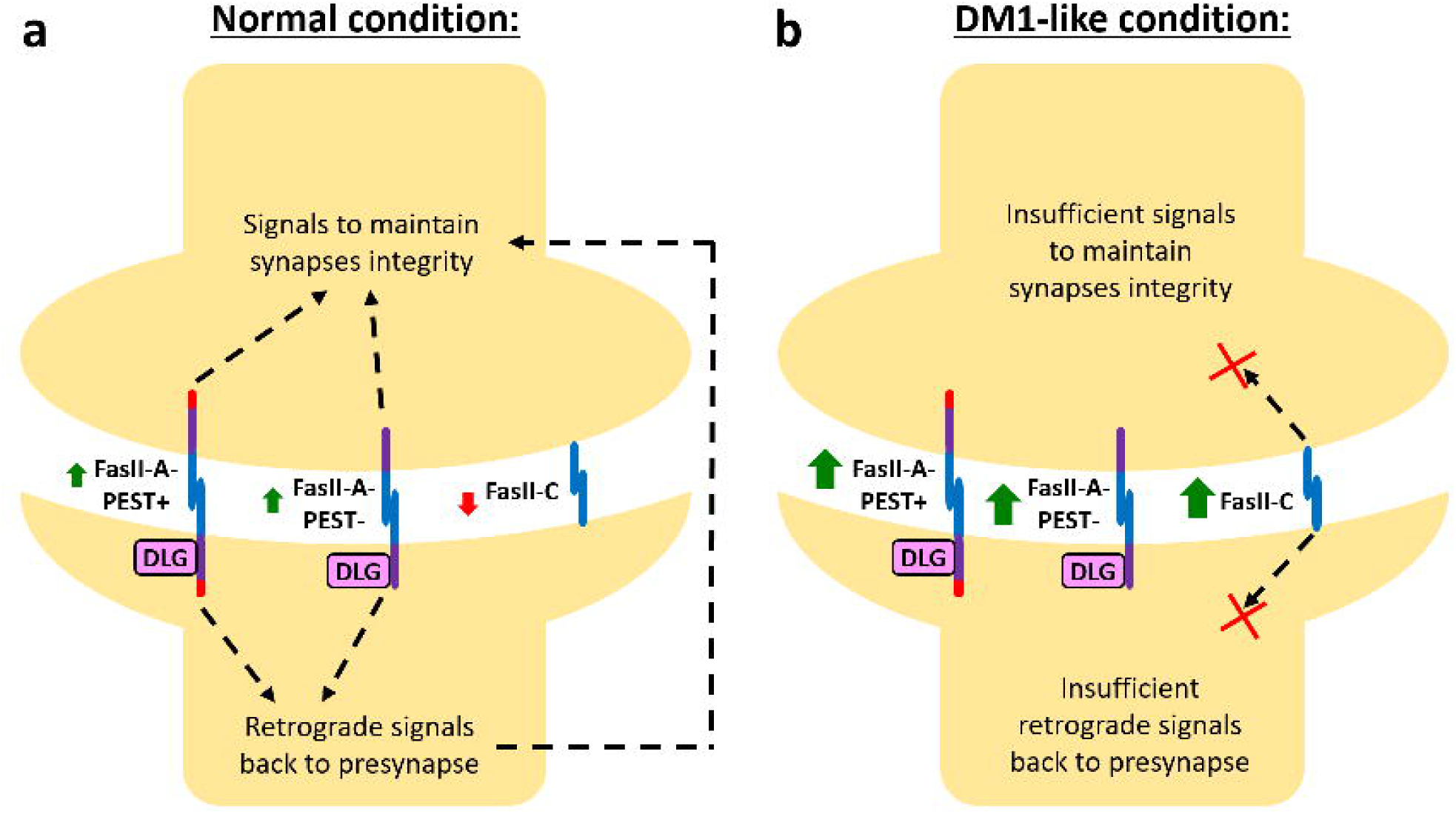
Working model of synaptopathology in a *Drosophila* larval NMJ model of DM1. **a** Under the normal condition, FasII-A-PEST+ and FasII-A-PEST-are the major isoforms present at the pre- and postsynaptic terminals. Both FasII-A isoforms convey intracellular signals to maintain synapse integrity. Postsynaptic FasII-A are capable of interacting with DLG, and may facilitate retrograde signals, such as Gbb (*Drosophila* counterpart of mammalian BMP), that help with maintaining bouton numbers. **b** Under the DM1-like condition, expanded *CUG* repeats causes upregulation of overall FasII, which in turn causes an abnormally high level of FasII-C at the synapse. FasII-C may bind to itself or compete with other isoforms to bind to the FasII-A isoforms. Since the FasII-C isoform lacks the cytoplasmic domain, its binding to FasII-A disrupts the intracellular signals required for maintaining synapse integrity.

It is likely that the intricate mechanisms of synapse regulation involving NCAM1 in mammals are far more complex than what we found in *Drosophila*. Nevertheless, our study provided the important foundation of a basic mechanism of synapse dysregulation in a simple DM1 model. It is possible that specific levels of different mammalian NCAM1 isoforms are expressed in different types of neurons and muscles for their proper functions. The expanded *CUG* RNA in DM1 likely disrupts the delicate regulation of NCAM1 and/or other cell adhesion molecules, which in turn causes synaptic dysfunction. Although we cannot directly change the genetic make-up of DM1 patients, we may be able to alleviate their CNS abnormalities by restoring the proper ratios of NCAM1 and/or other cell adhesion molecules at the synapses.

## Methods

### Fly Stocks

The following stocks were used: *w^1118^*, *elav^GS^-GAL4*, and *UAS-CD8::GFP* were acquired from Bloomington *Drosophila* Stock Center. *UAS-CUG_60_* and *UAS-CUG_480_* ^27^ were obtained from Ruben Artero. *UAS-CAG_250_*^49^ was from Nancy Bonini. *C380-GAL4* (a.k.a. *BG380*) ^50^ and *C57-GAL4* (a.k.a. *BG57*) ^50^ were obtained from Vivian Budnik. *UAS-FasII-A-PEST+* ^32, 33^, *UAS-FasII-A-PEST-* ^51^, *UAS-FasII-C* ^51^, *UAS-Total-FasII-RNAi* (a.k.a. *UAS-Total-FasII-dsRNA* or *P[KK100888]VIE-260B* ^40^, Vienna *Drosophila* RNAi Centre, #v103807) and *UAS-FasII-A-RNAi* ^39^ and *UAS-FasII-A&C-RNAi* (unpublished) were obtained from Brian McCabe. *UAS-FasII-A&C-RNAi* was originally named *UAS-FasII-C-RNAi(#38)* since it was designed against Exon 8 of *fasII*, which encodes for the GPI anchor of FasII-C. However, the construct was found to knock down FasII-A-PEST+ and FasII-A-PEST-as well (Supplementary Fig. 1 and Supplementary Fig. 2). Hence, it was renamed as *UAS-FasII-A&C-RNAi* in this study. *LexAop-FasII-A&C-RNAi* was generated by subcloning the same RNAi construct (Amplicon sequence: GCTAATAACA ATCTCGGCAC GTTGCTCTAT TCGGCCGGAT TTAATTCCGG TGTCGGTGCG CTACACAAAC GACTGTTCAC AACAACAACA ACAACAACAG CCACATCAAC AACAACAATC ACATCGATAA CAACAGCAAC AACAACAATC ATTACGCTGG CCAC) into the pJFRC19-13XLexAop2-IVS-myr::GFP vector (Addgene plasmid #26224), and inserted onto the 2^nd^ chromosome using the Phi3C1 system. *nSyb-LexA* (a.k.a. *nSyb-LexA-GAD*) ^52^ was obtained from Ching-Po Yang and Tzumin Lee. *Cass^M^-LexA* (unpublished) was obtained from Vivian Budnik. Flies were reared in standard *Drosophila* medium at 25°C. For genotypes involving *elav^GS^-GAL4*, RU486 was dissolved in 100% ethanol before added to the medium to achieve a final concentration of 50 µM.

### Semi-quantitative RT-PCR

mRNA was extracted from either dissected larval central nervous systems or body wall muscles using standard TRIzol extraction method as previously described ^53^. Synthesis of cDNA was performed using the ImProm-II^TM^ Reverse Transcription System (Promega). Primers used were as follow: Total-FasII Forward: GCAACCAGGTGGGATTAGG (on Exon 6), Total-FasII Reverse: TAACGCCCGGACAGTATTTG (partially on Exon 6 and partially on Exon 7), FasII-A Forward: CTGTCCGGGCGTTAAGATC (partially on Exon 6 and partially on Exon 7), FasII-A Reverse: ACGTCAATTCCTCGTGTCG (partially on Exon 7 and partially on TM domain), FasII-A-PEST+ Forward: ACACGAGGAATTGACGTCATC (partially on Exon 7 and partially on TM domain), FasII-A-PEST+ Reverse: GTGGCTCCTTTACCAGCTG (on the PEST domain), FasII-C Forward: GCGTTAAGATCAGCGGCAC (on Exon 7), FasII-C Reverse: GAATCGGACTCACCTCGTG (partially on Exon 7 and partially on GPI anchor). The designs of these primers had been previously described ^39^.

### Locomotor behavioral assay for larvae

Larval crawling assay was previously described ^54^. In brief, wandering third instar larvae were loaded onto a 2% agarose plate with a grid (0.5 cm^2^ per square) placed underneath. The total number of gridlines crossed by the animals in one minute was counted, and the actual distance calculated. *n* represents the number of larvae analyzed.

### Immunohistochemistry

Larval body-wall muscles were dissected and fixed for 10 min in 4% paraformaldehyde, and permeabilized with 0.2% Triton-X 100 at room temperature. Details were previously described in ^55^. Antibodies and their concentrations used: anti-HRP-Alexa Fluor 488 1:500 (Jackson), anti-HRP-Alex Fluor 647 1:500 (Jackson), anti-DLG1 1:750 (4F3, DSHB), anti-FasII 1:50 (1D4, DSHB). Secondary antibodies conjugated to Alexa Fluor 488/594 (Abcam) were used at a concentration of 1:200.

### Quantification of boutons

The number of type I boutons was obtained at muscles 6 and 7 of abdominal segment A3 of late 3^rd^ instar larvae unless specified otherwise. *n* represents the number of analyzed NMJs. At most two NMJs were quantified in each animal. Details were previously described in ^55, 56^. The development of larvae expressing *CUG_480_* using *C380* alone, *C380+C57* or *elav^GS^* (fed with food with RU486) were slower than that of controls by approximately 24 hours. Thus, crosses involving these genotypes were set up one day in advance so the dissections and immunostaining can be carried out at the same time.

### Quantification of muscle areas

Lengths and widths of muscle 6 of segment A3 were measured under a 20x objective on a widefield microscope, and the areas were subsequently calculated (Supplementary Fig. 6). *N* represents the number of analyzed muscles. At most two muscles were quantified in each animal.

### Slot blot assay for *Drosophila* FasII and mice NCAM1

Fly heads, fly thorax muscles, mouse CNS tissues, or mouse muscle tissues were homogenized in the lysis buffer (100 mM Tris/HCl, pH6.8; 2% Sodium Dodecyl Sulfate; 40% w/v Glycerol). The homogenates were boiled at 99°C for 10 min, then centrifuged at 14,000 x *g* for 2 min and the supernatant was collected. Before the slot blot assay, samples were diluted with 2% SDS to a final volume of 200 µL and heated at 99°C for 10 min. Protein samples were loaded to a 48-well Bio-Dot® microfiltration apparatus (Bio-Rad Laboratories, Hercules, CA, USA) with BioTrace^TM^ NT nitrocellulose membrane (Pall Life Sciences, Portsmouth, UK; pore size 0.2 µm). The membrane was blocked at room temperature in 5% non-fat milk and incubated in 34B3 (Developmental Studies Hybridoma Bank, Iowa City, IA, USA; Mouse; 1:10) at 4°C overnight for the detection of FasII in fly protein samples. For mouse protein samples, the membrane was incubated in ab154566 (Abcam, Cambridge, UK; Rabbit; 1:1,000) for NCAM1 detection. The protein chemiluminescence signal was obtained and visualized with the ChemiDoc^TM^ Touch Gel Imaging System (Bio-Rad Laboratories, Hercules, CA, USA). After FasII or NCAM1 detection, the membrane was stripped in stripping buffer (Thermo Fisher Scientific, Grand Island, NY, USA) and blocked again in 5% non-fat milk. β-tubulin was used as the internal loading control, where the membrane was re-probed with ab6046 (Abcam, Cambridge, UK; Rabbit; 1:500) for fly protein samples or 2G7D4 (GenScript, Piscataway, NJ, USA; Mouse; 1:2,000) for mouse protein samples, respectively. The images were analyzed with ImageLab^TM^ software (Bio-Rad Laboratories, Hercules, CA, USA) and band intensities were quantified using the Image J software (Research Services Branch, National Institute of Mental Health).

### Slot blot assay for human frontal cortex

Total protein was extracted from 20-30 mg human frontal cortex using RIPA buffer (Pierce^TM^ RIPA Buffer, ThermoScientific, 89901) supplemented with 0.05% CHAPS (Sigma, C3023), 1X complete protease inhibitor (Sigma-Aldrich, 04693124001), 1X PhosSTOP phosphatase inhibitor (Sigma-Aldrich, 04906845001) and 1 mM sodium orthovanadate (Sigma, S6508). Protein concentrations were determined using the Pierce BCA Protein Assay Kit (Thermo Scientific, 23227). Porablot NCP nitrocellulose membranes (Macherey-Nagel, 741280) and Bio-Dot® SF filter paper (Bio-Rad, 1620161) were soaked and equilibrated in 1X PBS (Gibco, 20012-019) and then placed in the Bio-Dot® SF Apparatus (Bio-Rad, 1706542). Membrane slots were rinsed with 100 µL 1X PBS, and then loaded with 20 µg frontal cortex protein in 100 µL RIPA buffer under vacuum. The membranes were carefully removed from the Bio-Dot® SF Apparatus and stained with ATX Ponceau S red solution (Sigma-Aldrich, 09189-1L-F) for total protein quantification. Membranes were blocked in 5% Blotto (Santa Cruz Biotech, sc-2324) diluted in 1X TBS-T (10 mM Tris-HCl, 0.15 M NaCl, 0.05% Tween 20) for 1 hour at room temperature, and incubated overnight at 4°C with primary anti-NCAM1 antibody (GeneTex, GTX111684) diluted 1:10,000 in 5% Blotto. After three washes in 1X TBS-T, membranes were incubated with secondary HRP-conjugated goat anti-rabbit IgG (LifeTechnologies, 31460) diluted 1:5,000 in 5% Blotto, for 1h at room temperature. Membranes were finally washed another three times in 1X TBS-T and developed with Clarity Max ECL Substrate (BioRad, 1705062). Image acquisition and quantification were conducted with BioRad ChemiDoc^TM^ MP Imaging System and software (Image Lab 6.0.1). Relative protein levels in non-DM controls were normalized to 1 and statistical analysis was performed with Prism (GraphPad Software, Inc).

### Transdifferentiation of myotubes

Control or DM1 patient myoblasts were differentiated for 7 days into transdifferentiated myotubes. Differentiation conditions were described in ^57^.

### Quantitative dot blot assay

1 μg/well of protein samples were denatured (100°C for 5 min) and loaded in quantitative dot blot (QDB) plates (Quanticision Diagnostics Inc, Research Triangle Park, NC, USA). Each cell sample was loaded in quadruplicate in two different plates; one was used to detect NCAM11 (1:1,000, proteintech, Manchester, UK,) and the other for GAPDH (1:500, (G-9) Santa Cruz Biotechnology, Dallas, TX, USA), which was used as an endogenous control. Details were previously described in ^58^.

### Electrophysiology

Electrophysiology was performed as previously described ^59^. For detailed methods on how to perform Drosophila NMJ electrophysiology, please see published protocols ^60, 61^. Briefly, wandering third instar larvae were collected and filleted for NMJ analysis. Driver control and experimental electrophysiological recordings were performed in parallel using identical conditions. Larval dissections and recordings were performed in a modified, low-magnesium HL3 saline ^62^. (70 mM NaCl, 5 mM KCl, 5mM HEPES, 10 mM NaHCO_3_, 115 mM sucrose, 4.2 mM trehalose, 0.5 mM CaCl_2_ (unless otherwise noted), 10 mM MgCl_2_, pH 7.2. Neuromuscular junction sharp electrode recordings were performed on muscles 6/7 of abdominal segments A2 or A3. For a muscle to be acceptable for recording, it needed to have an input resistance of ≥ 4 MΩ and a resting potential more hyperpolarized than −58 mV.

Recordings were performed on an Olympus BX51WI microscope and acquired using an Axoclamp 900A amplifier, Digidata 1440A acquisition system and pClamp10.7 (Molecular Devices) software. Data were analyzed using MiniAnalysis (Synaptosoft) and the Clampfit (Molecular Devices) programs. Miniature excitatory postsynaptic potentials (mEPSPs) and excitatory postsynaptic potentials (EPSPs at 1 Hz stimulus) were collected as previously described ^59^. For each NMJ recorded, four quantifications were made: quantal size (mEPSP size); quantal frequency (mEPSP frequency); evoked potential size (EPSP size); and quantal content (QC). In the case of QC, uncorrected QC was calculated per NMJ by dividing the average EPSP by the average mEPSP.

### Statistical analysis

For comparisons between three or more sample groups, an analysis of variance (ANOVA) with Tukey post-hoc test was performed. For pair-wise comparisons a Student *t*-test was used. \**P*<0.05, ** *P*<0.01, *** *P*<0.001, **** *P*<0.0001. All histograms depict mean ± S.E.M. All experiments were performed at least three times independently.

## Acknowledgements

This work was supported by Research Grants Council of Hong Kong (14102220), Jérôme Lejeune Foundation (GRT-2022A / 2120) and AFM-Téléthon. We would also like to thank Dr. Ching-Po Yang and Prof. Tzumin Lee (Life Sciences Institute, University of Michigan, USA) for sharing of *Drosophila* reagents.

## Author contributions

A.C.K. designed and performed most experiments and contributed to manuscript writing; K.Y.W.Y., L.I.L., Z.S.C. and J.M.S.F. contributed to various experiments, including RT-PCRs, slot blot assays and dissections for NMJ bouton analyses; S.I.P. and Y.W. conducted larval behavioral assays; N.S.A. and C.A.F. contributed to electrophysiology experiments and analyses; A.B. and R.A. provided human DM1 patient samples and patient cell-derived samples, and performed quantitative dot blot analyses; P.M. and M.G. provided human DM1 patient frontal cortex samples and performed slot blot analyses; A.H. and G.G. provided DMSXL mice samples; C.K.B. and V.B. generated the *Cass^M^-LexA* fly line and provided numerous key fly lines and antibodies for the study; E.S.B. and B.M. generated the *UAS-FasII-A&C-RNAi* fly line and provided numerous key fly lines for the study; K.Y.W.Y., L.I.L., A.B., M.G. and C.A.F. contributed to manuscript writing; H.Y.E.C. directed the project and contributed to manuscript writing.

## References

1. Johnson NE, et al. Population-Based Prevalence of Myotonic Dystrophy Type 1 Using Genetic Analysis of Statewide Blood Screening Program. Neurology 96, e1045–e1053 (2021).

2. Harper PS, Harper PS. Myotonic dystrophy, 3rd edn. W.B. Saunders (2001).

3. Patel N, et al. Neurobehavioral Phenotype of Children With Congenital Myotonic Dystrophy. Neurology 102, e208115 (2024).

4. Brook JD, et al. Molecular basis of myotonic dystrophy: expansion of a trinucleotide (CTG) repeat at the 3’ end of a transcript encoding a protein kinase family member. Cell 69, 385 (1992).

5. Fu YH, et al. An unstable triplet repeat in a gene related to myotonic muscular dystrophy. Science 255, 1256–1258 (1992).

6. Mahadevan M, et al. Myotonic dystrophy mutation: an unstable CTG repeat in the 3’ untranslated region of the gene. Science 255, 1253–1255 (1992).

7. Thornton CA, Johnson K, Moxley RT, 3rd. Myotonic dystrophy patients have larger CTG expansions in skeletal muscle than in leukocytes. Ann Neurol 35, 104–107 (1994).

8. Mankodi A, et al. Myotonic dystrophy in transgenic mice expressing an expanded CUG repeat. Science 289, 1769–1773 (2000).

9. Gladman JT, Mandal M, Srinivasan V, Mahadevan MS. Age of onset of RNA toxicity influences phenotypic severity: evidence from an inducible mouse model of myotonic dystrophy (DM1). PLoS One 8, e72907 (2013).

10. Mankodi A, et al. Muscleblind localizes to nuclear foci of aberrant RNA in myotonic dystrophy types 1 and 2. Hum Mol Genet 10, 2165–2170 (2001).

11. Ho TH, Charlet BN, Poulos MG, Singh G, Swanson MS, Cooper TA. Muscleblind proteins regulate alternative splicing. EMBO J 23, 3103–3112 (2004).

12. Jiang H, Mankodi A, Swanson MS, Moxley RT, Thornton CA. Myotonic dystrophy type 1 is associated with nuclear foci of mutant RNA, sequestration of muscleblind proteins and deregulated alternative splicing in neurons. Hum Mol Genet 13, 3079–3088 (2004).

13. Koon AC, Chan HY. Drosophila melanogaster As a Model Organism to Study RNA Toxicity of Repeat Expansion-Associated Neurodegenerative and Neuromuscular Diseases. Front Cell Neurosci 11, 70 (2017).

14. Gourdon G, Meola G. Myotonic Dystrophies: State of the Art of New Therapeutic Developments for the CNS. Front Cell Neurosci 11, 101 (2017).

15. Hernandez-Hernandez O, et al. Myotonic dystrophy CTG expansion affects synaptic vesicle proteins, neurotransmission and mouse behaviour. Brain 136, 957–970 (2013).

16. Marteyn A, et al. Mutant human embryonic stem cells reveal neurite and synapse formation defects in type 1 myotonic dystrophy. Cell Stem Cell 8, 434–444 (2011).

17. Angeard N, Gargiulo M, Jacquette A, Radvanyi H, Eymard B, Heron D. Cognitive profile in childhood myotonic dystrophy type 1: is there a global impairment? Neuromuscul Disord 17, 451–458 (2007).

18. Angeard N, et al. A new window on neurocognitive dysfunction in the childhood form of myotonic dystrophy type 1 (DM1). Neuromuscul Disord 21, 468–476 (2011).

19. Douniol M, et al. Psychiatric and cognitive phenotype of childhood myotonic dystrophy type 1. Dev Med Child Neurol 54, 905–911 (2012).

20. Douniol M, et al. Psychiatric and cognitive phenotype in children and adolescents with myotonic dystrophy. Eur Child Adolesc Psychiatry 18, 705–715 (2009).

21. Ekstrom AB, Hakenas-Plate L, Samuelsson L, Tulinius M, Wentz E. Autism spectrum conditions in myotonic dystrophy type 1: a study on 57 individuals with congenital and childhood forms. Am J Med Genet B Neuropsychiatr Genet 147B, 918–926 (2008).

22. Meola G, Sansone V. Cerebral involvement in myotonic dystrophies. Muscle Nerve 36, 294–306 (2007).

23. Peric S, et al. Peripheral neuropathy in patients with myotonic dystrophy type 1. Neurol Res 35, 331–335 (2013).

24. Gantelet E, Kraftsik R, Delaloye S, Gourdon G, Kuntzer T, Barakat-Walter I. The expansion of 300 CTG repeats in myotonic dystrophy transgenic mice does not induce sensory or motor neuropathy. Acta Neuropathol 114, 175–185 (2007).

25. Panaite PA, Kielar M, Kraftsik R, Gourdon G, Kuntzer T, Barakat-Walter I. Peripheral neuropathy is linked to a severe form of myotonic dystrophy in transgenic mice. J Neuropathol Exp Neurol 70, 678–685 (2011).

26. Ruiz-Canada C, Budnik V. Introduction on the use of the Drosophila embryonic/larval neuromuscular junction as a model system to study synapse development and function, and a brief summary of pathfinding and target recognition. Int Rev Neurobiol 75, 1–31 (2006).

27. Garcia-Lopez A, Monferrer L, Garcia-Alcover I, Vicente-Crespo M, Alvarez-Abril MC, Artero RD. Genetic and chemical modifiers of a CUG toxicity model in Drosophila. PLoS One 3, e1595 (2008).

28. Chen K, Featherstone DE. Discs-large (DLG) is clustered by presynaptic innervation and regulates postsynaptic glutamate receptor subunit composition in Drosophila. BMC Biol 3, 1 (2005).

29. Lahey T, Gorczyca M, Jia XX, Budnik V. The Drosophila tumor suppressor gene dlg is required for normal synaptic bouton structure. Neuron 13, 823–835 (1994).

30. Picchio L, Plantie E, Renaud Y, Poovthumkadavil P, Jagla K. Novel Drosophila model of myotonic dystrophy type 1: phenotypic characterization and genome-wide view of altered gene expression. Hum Mol Genet 22, 2795–2810 (2013).

31. Ashley J, Packard M, Ataman B, Budnik V. Fasciclin II signals new synapse formation through amyloid precursor protein and the scaffolding protein dX11/Mint. J Neurosci 25, 5943–5955 (2005).

32. Schuster CM, Davis GW, Fetter RD, Goodman CS. Genetic dissection of structural and functional components of synaptic plasticity. II. Fasciclin II controls presynaptic structural plasticity. Neuron 17, 655–667 (1996).

33. Schuster CM, Davis GW, Fetter RD, Goodman CS. Genetic dissection of structural and functional components of synaptic plasticity. I. Fasciclin II controls synaptic stabilization and growth. Neuron 17, 641–654 (1996).

34. Osterwalder T, Yoon KS, White BH, Keshishian H. A conditional tissue-specific transgene expression system using inducible GAL4. Proc Natl Acad Sci U S A 98, 12596–12601 (2001).

35. Gomes-Pereira M, et al. CTG trinucleotide repeat “big jumps“: large expansions, small mice. PLoS Genet 3, e52 (2007).

36. Wright JW, Copenhaver PF. Different isoforms of fasciclin II play distinct roles in the guidance of neuronal migration during insect embryogenesis. Dev Biol 225, 59–78 (2000).

37. Grenningloh G, Rehm EJ, Goodman CS. Genetic analysis of growth cone guidance in Drosophila: fasciclin II functions as a neuronal recognition molecule. Cell 67, 45–57 (1991).

38. Fankhauser N, Maser P. Identification of GPI anchor attachment signals by a Kohonen self-organizing map. Bioinformatics 21, 1846–1852 (2005).

39. Beck ES, et al. Regulation of Fasciclin II and synaptic terminal development by the splicing factor beag. J Neurosci 32, 7058–7073 (2012).

40. Dietzl G, et al. A genome-wide transgenic RNAi library for conditional gene inactivation in Drosophila. Nature 448, 151–156 (2007).

41. Lai SL, Lee T. Genetic mosaic with dual binary transcriptional systems in Drosophila. Nat Neurosci 9, 703–709 (2006).

42. Thomas U, et al. Synaptic clustering of the cell adhesion molecule fasciclin II by discs-large and its role in the regulation of presynaptic structure. Neuron 19, 787–799 (1997).

43. Zito K, Fetter RD, Goodman CS, Isacoff EY. Synaptic clustering of Fascilin II and Shaker: essential targeting sequences and role of Dlg. Neuron 19, 1007–1016 (1997).

44. Berke B, Wittnam J, McNeill E, Van Vactor DL, Keshishian H. Retrograde BMP signaling at the synapse: a permissive signal for synapse maturation and activity-dependent plasticity. J Neurosci 33, 17937–17950 (2013).

45. Neuert H, et al. The Drosophila NCAM homolog Fas2 signals independently of adhesion. Development 147, (2020).

46. Kristiansen LV, Hortsch M. Fasciclin II: the NCAM ortholog in Drosophila melanogaster. Adv Exp Med Biol 663, 387–401 (2010).

47. Santoro L, et al. Perioral skin biopsy to study skeletal muscle protein expression. Muscle Nerve 41, 392–398 (2010).

48. Otero BA, et al. Transcriptome alterations in myotonic dystrophy frontal cortex. Cell Rep 34, 108634 (2021).

49. Li LB, Yu Z, Teng X, Bonini NM. RNA toxicity is a component of ataxin-3 degeneration in Drosophila. Nature 453, 1107–1111 (2008).

50. Budnik V, et al. Regulation of synapse structure and function by the Drosophila tumor suppressor gene dlg. Neuron 17, 627–640 (1996).

51. Lin DM, Goodman CS. Ectopic and increased expression of Fasciclin II alters motoneuron growth cone guidance. Neuron 13, 507–523 (1994).

52. Pfeiffer BD, et al. Refinement of tools for targeted gene expression in Drosophila. Genetics 186, 735–755 (2010).

53. Li L, Ng NK, Koon AC, Chan HY. Expanded polyalanine tracts function as nuclear export signals and promote protein mislocalization via eEF1A1 factor. J Biol Chem 292, 5784–5800 (2017).

54. Bai Y, et al. Integrating Display and Delivery Functionality with a Cell Penetrating Peptide Mimic as a Scaffold for Intracellular Multivalent Multitargeting. J Am Chem Soc 138, 9498–9507 (2016).

55. Koon AC, et al. Drosophila Exo70 Is Essential for Neurite Extension and Survival under Thermal Stress. J Neurosci 38, 8071–8086 (2018).

56. Koon AC, et al. Autoregulatory and paracrine control of synaptic and behavioral plasticity by octopaminergic signaling. Nat Neurosci 14, 190–199 (2011).

57. Arandel L, et al. Immortalized human myotonic dystrophy muscle cell lines to assess therapeutic compounds. Dis Model Mech 10, 487–497 (2017).

58. Moreno N, Gonzalez-Martinez I, Artero R, Cerro-Herreros E. Rapid Determination of MBNL1 Protein Levels by Quantitative Dot Blot for the Evaluation of Antisense Oligonucleotides in Myotonic Dystrophy Myoblasts. Methods Mol Biol 2434, 207–215 (2022).

59. Armstrong NS, Frank CA. The calcineurin regulator Sarah enables distinct forms of homeostatic plasticity at the Drosophila neuromuscular junction. Front Synaptic Neurosci 14, 1033743 (2022).

60. Imlach W, McCabe BD. Electrophysiological methods for recording synaptic potentials from the NMJ of Drosophila larvae. J Vis Exp, (2009).

61. Zhang B, Stewart B. Synaptic Electrophysiology of the Drosophila Neuromuscular Junction. Cold Spring Harb Protoc, (2024).

62. Stewart BA, Atwood HL, Renger JJ, Wang J, Wu CF. Improved stability of Drosophila larval neuromuscular preparations in haemolymph-like physiological solutions. J Comp Physiol A 175, 179–191 (1994).

