## Supplementary Information for "Upregulation of FasII underlies synergistic neuropathological and behavioral defects in a *Drosophila* model of myotonic dystrophy"

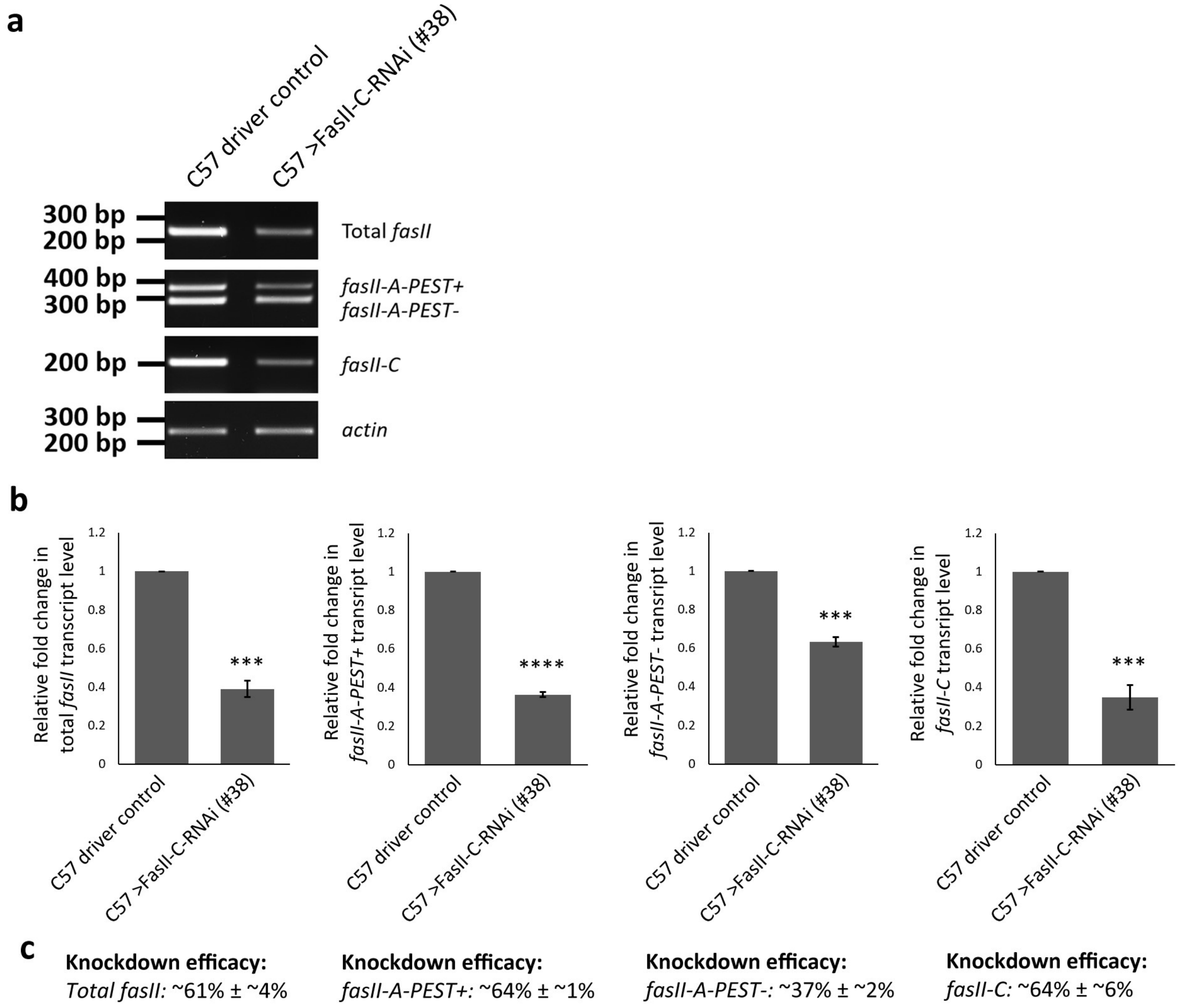


**Supplementary Figure 1.** ***UAS-FasII-C (#38)*** **knocks down at least three isoforms of** ***fasII*****.** **(a)** Representative semi-quantitative RT-PCR of total *fasII* and various *fasII* isoforms in *Drosophila* larval body wall muscles (BWMs). *C57-Gal4* is a BWM driver. **(b)** Quantification of **(a)**. **(c)** Knockdown efficacies of total *fasII* and various *fasII* isoforms. $N = 3$. Histograms depicts mean ± SEM. ***$p < 0.001$, ****$p < 0.0001$.

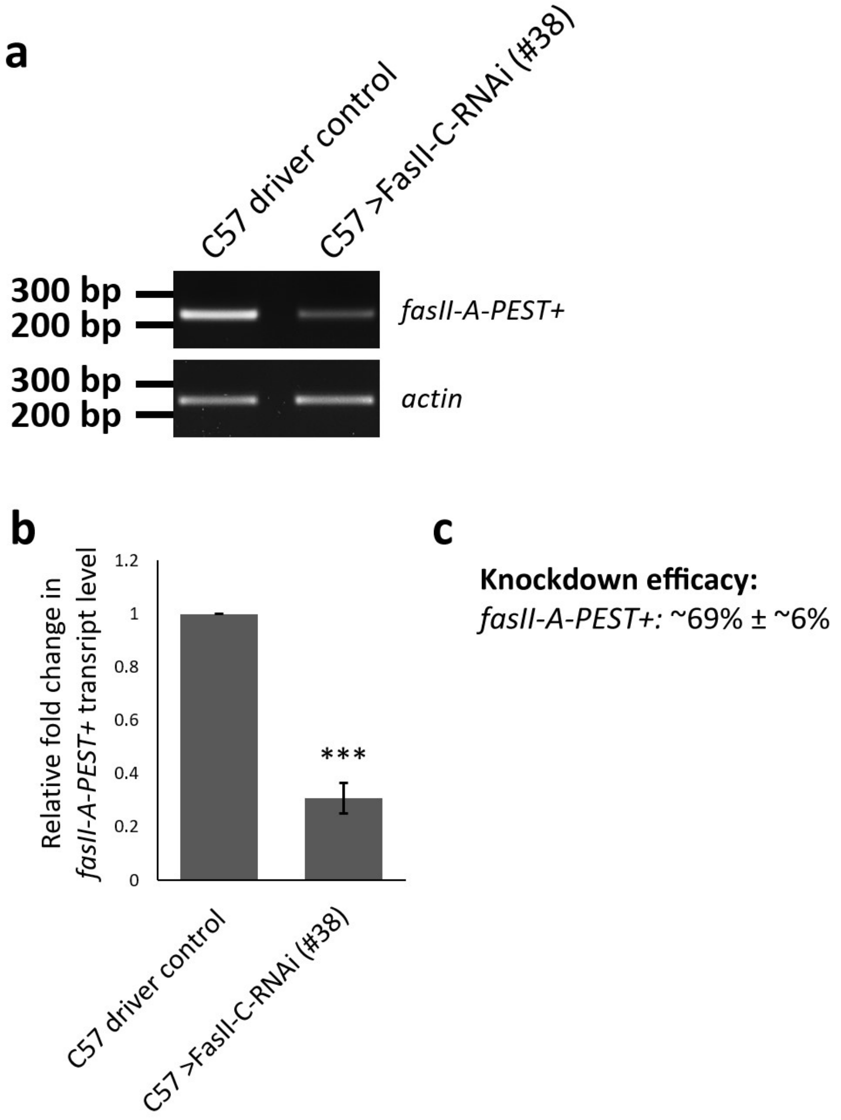


**Supplementary Figure 2.** ***UAS-FasII-C (#38)*** **knocks down the PEST+ isoform of** ***fasII*****.** **(a)** Representative semi-quantitative RT-PCR of *fasII-A-PEST+* in *Drosophila* larval BWMs. The forward primer used in this PCR targets the nucleotide sequence of Exon 7 and the transmembrane domain, while the reverse primer targets the nucleotide sequence of the PEST domain. **(b)** Quantification of **(a)**. $N = 3$. **(c)** Knockdown efficacy of *fasII-A-PEST+*. Histograms depicts mean ± SEM. ***$p < 0.001$.

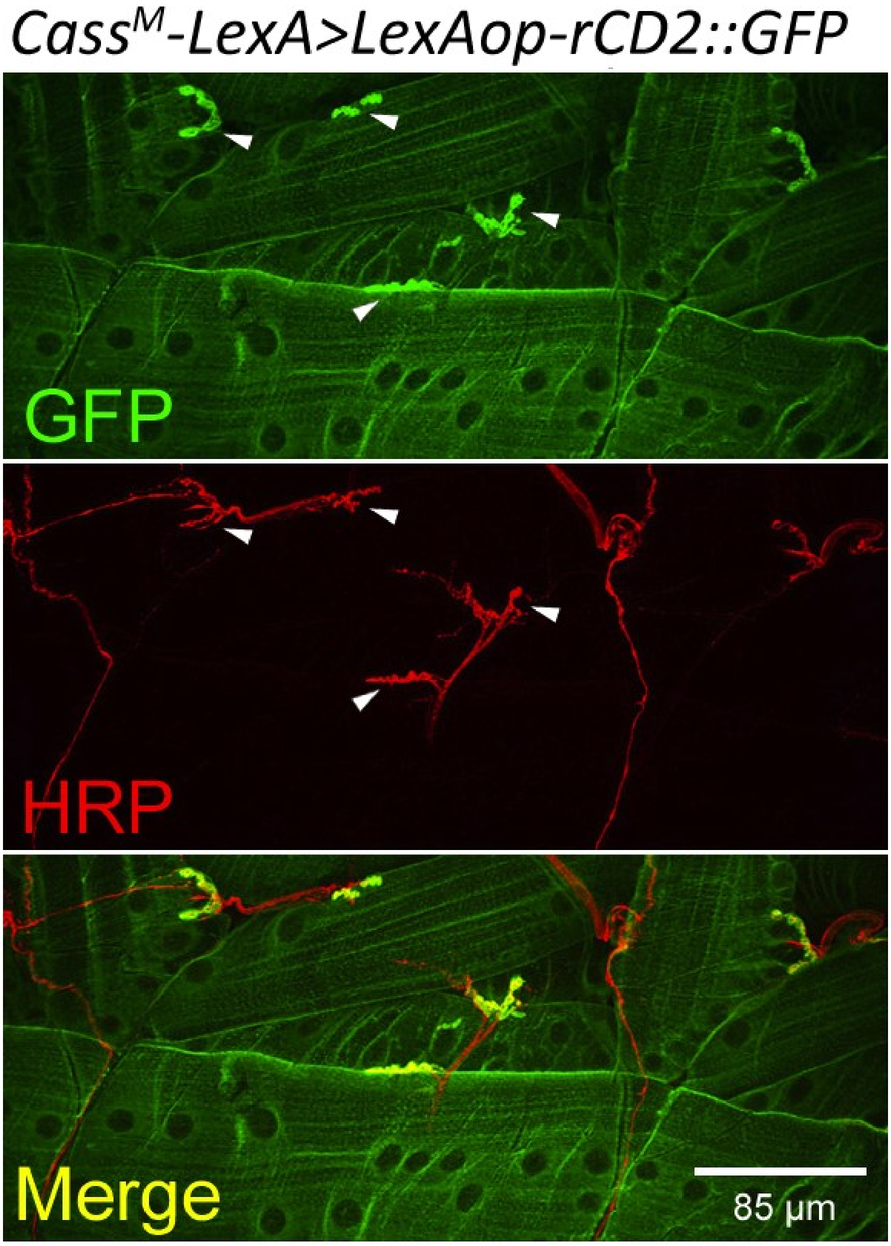


**Supplementary Figure 3.** ***Cass**[M]**-LexA*** **expresses in body wall muscles but not in motor neurons at the *Drosophila* larval NMJ.** Confocal micrograph of *Drosophila* NMJs of late 3rd instar larvae on muscle 5, 6, 8, 12 and 13 of segment A3. Anti-GFP (in green) marks the membrane of the body wall muscles. Anti-HRP (in red) marks the motor neurons. White arrowheads denote regions of synaptic boutons. On the muscles, the membrane-rich subsynaptic reticulum (SSR) surrounds the boutons. Thus, strong green signals are observed at the SSR, indicating abundant amounts of membrane-tethered rCD2::GFP. Scale bar is 85 µm.

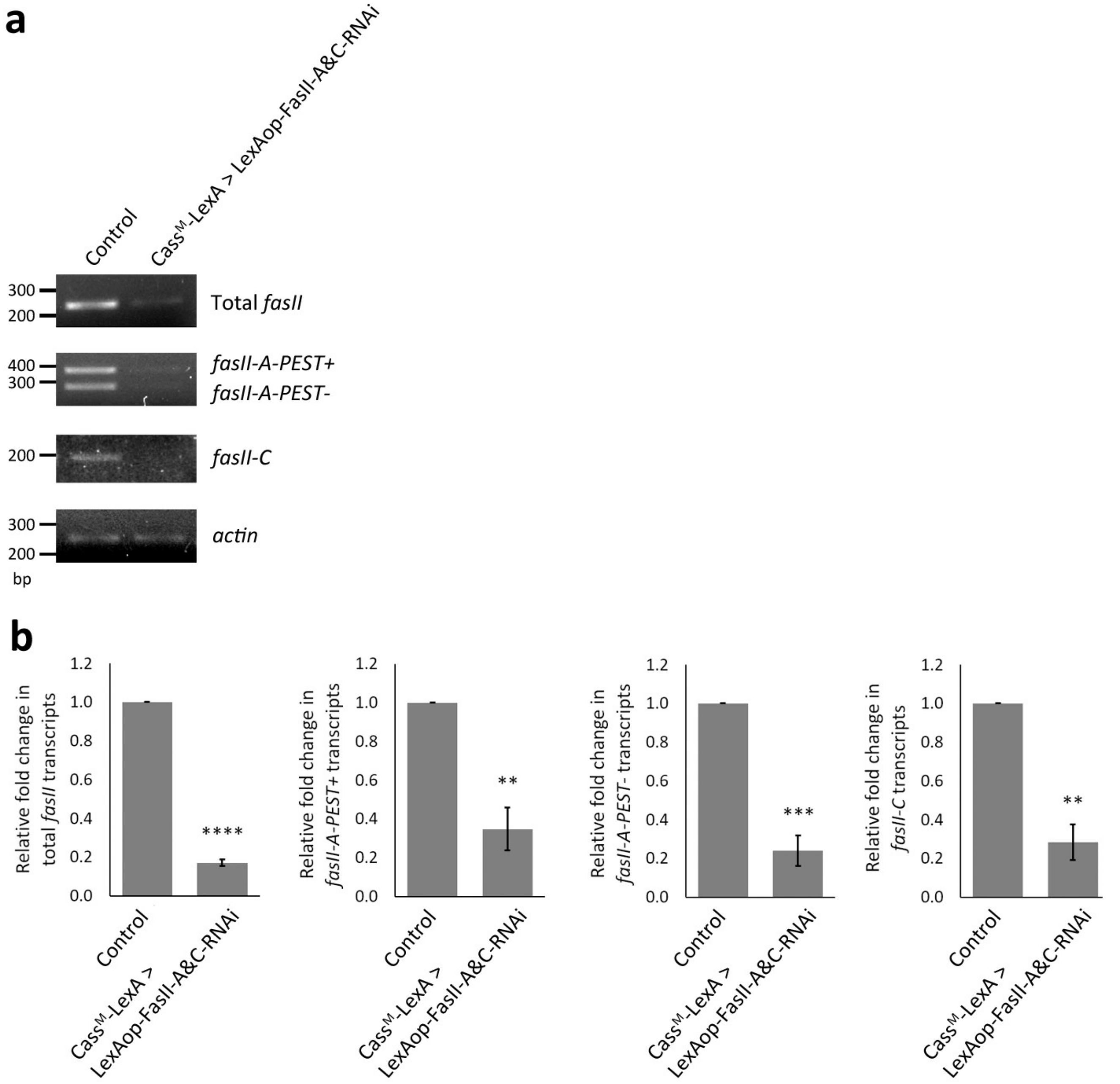


**c**

**Knockdown efficacy:**
*Total fasII:* ~83% ± ~2%

**Knockdown efficacy:**
*fasII-A-PEST+:* ~65% ± ~11%

**Knockdown efficacy:**
*fasII-A-PEST-:* ~76% ± ~8%

**Knockdown efficacy:**
*fasII-C:* ~72% ± ~9%

**Supplementary Figure 4. *LexAop-FasII-A&C-RNAi* knocks down the *fasII-A* and *fasII-C* isoforms. (a)** Representative semi-quantitative RT-PCR of total *fasII* and various *fasII* isoforms in *Drosophila* larval body wall muscles. **(b)** Quantification of **(a)**. **(c)** Knockdown efficacies of total *fasII* and various *fasII* isoforms. $N = 3$. Histograms depicts mean ± SEM. ***$p < 0.001$, ****$p < 0.0001$.

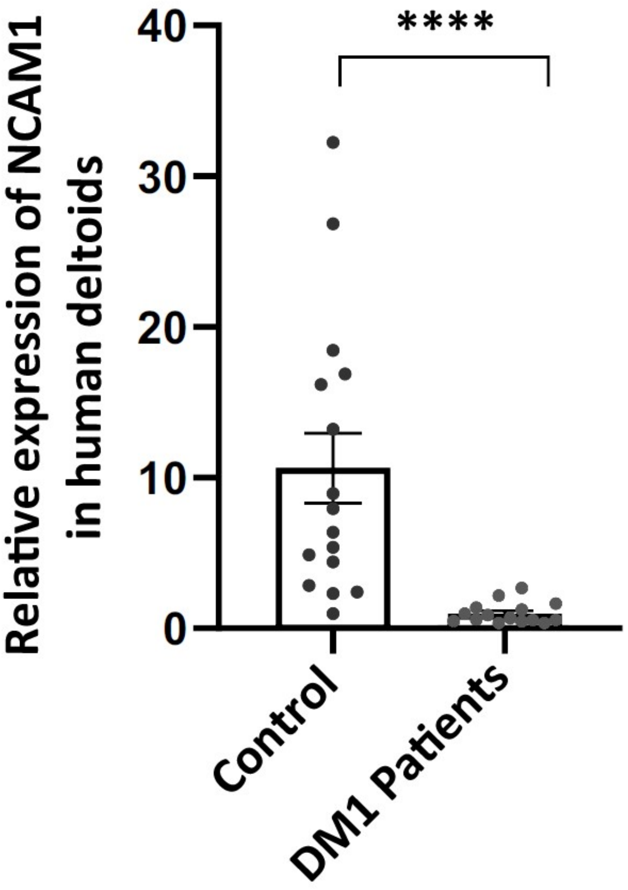


**Supplementary Figure 5. Lower expression of NCAM1 was detected in the deltoids of DM1 patients.** Quantitative dot blot data of NCAM1 in human deltoids. $n = 17$ for both control and DM1 patients. Histograms depicts mean ± SEM. ****$p < 0.0001$.

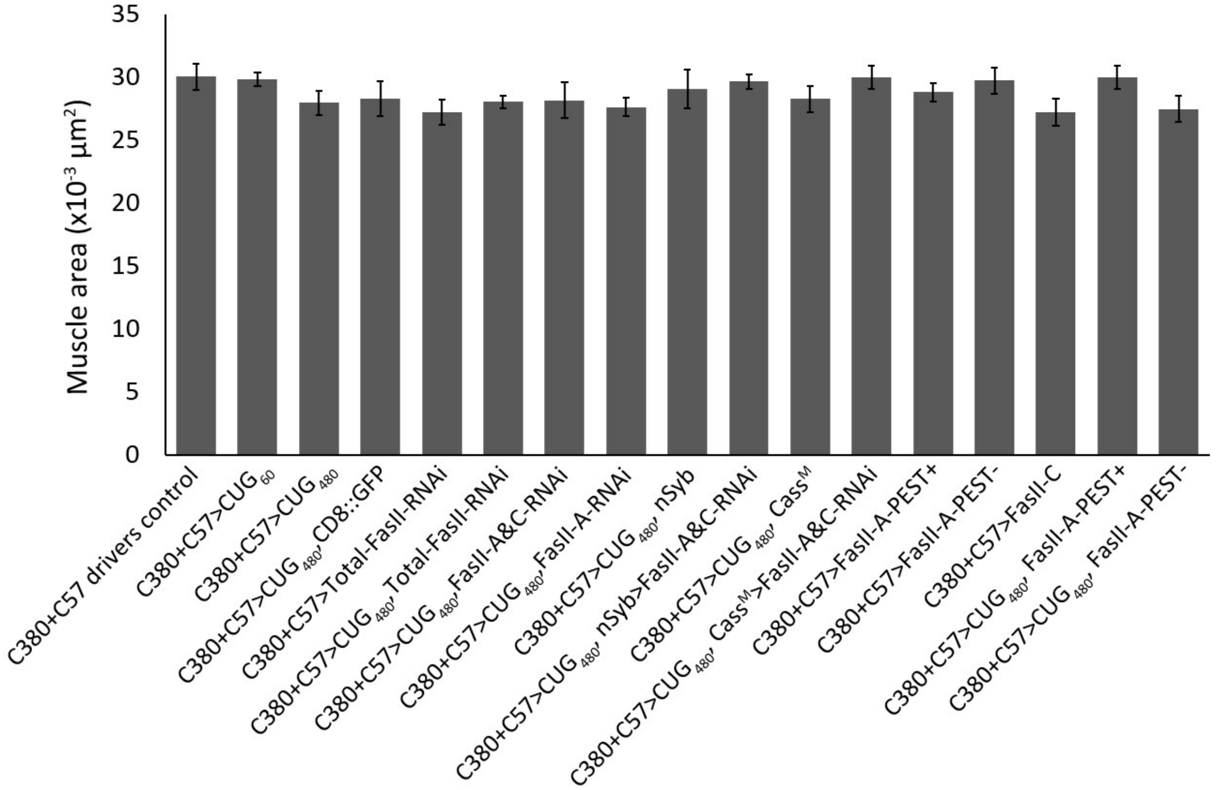


**Supplementary Figure 6. No notable difference was found among size of body wall muscles of the genotypes used in this study.** Quantification of muscle area (Muscle 6, Segment A3) for all genotypes used in this study. *n* = 10 for all genotypes, where *n* is the number of muscles analyzed.
